# Diaci v3.0: Chromosome-level assembly, *de novo* transcriptome and manual annotation of *Diaphorina citri,* insect vector of Huanglongbing

**DOI:** 10.1101/869685

**Authors:** Teresa D. Shippy, Prashant S. Hosmani, Mirella Flores-Gonzalez, Marina Mann, Sherry Miller, Matthew T. Weirauch, Chad Vosburg, Crissy Massimino, Will Tank, Lucas de Oliveira, Chang Chen, Stephanie Hoyt, Rebekah Adams, Samuel Adkins, Samuel T. Bailey, Xiaoting Chen, Nina Davis, Yesmarie DeLaFlor, Michelle Espino, Kylie Gervais, Rebecca Grace, Douglas Harper, Denisse L. Hasan, Maria Hoang, Rachel Holcomb, Margaryta R. Jernigan, Melissa Kemp, Bailey Kennedy, Kyle Kercher, Stefan Klaessan, Angela Kruse, Sophia Licata, Andrea Lu, Ron Masse, Anuja Mathew, Sarah Michels, Elizabeth Michels, Alan Neiman, Seantel Norman, Jordan Norus, Yasmin Ortiz, Naftali Panitz, Thomson Paris, Kitty M. R. Perentesis, Michael Perry, Max Reynolds, Madison M. Sena, Blessy Tamayo, Amanda Thate, Sara Vandervoort, Jessica Ventura, Nicholas Weis, Tanner Wise, Robert G. Shatters, Michelle Heck, Joshua B. Benoit, Wayne B. Hunter, Lukas A. Mueller, Susan J. Brown, Tom D’Elia, Surya Saha

## Abstract

**Background:** *Diaphorina citri* is a vector of “*Candidatus* Liberibacter asiaticus” (*C*Las), the gram-negative bacterial pathogen associated with citrus greening disease. Control measures rely on pesticides with negative impacts on the environment, natural ecosystems and human and animal health. In contrast, gene-targeting methods have the potential to specifically target the vector species and/or reduce pathogen transmission.

**Results:** To improve the genomic resources needed for targeted pest control, we assembled a *D. citri* genome based on PacBio long reads followed by proximity ligation-based scaffolding. The 474 Mb genome has 13 chromosomal-length scaffolds. 1,036 genes were manually curated as part of a community annotation project, composed primarily of undergraduate students. We also computationally identified a total of 1,015 putative transcription factors (TFs) and were able to infer motifs for 337 TFs (33 %). In addition, we produced a genome-independent transcriptome and genomes for *D. citri* endosymbionts.

**Conclusions:** Manual annotation provided more accurate gene models for use by researchers and also provided an excellent training opportunity for students from multiple institutions. All resources are available on CitrusGreening.org and NCBI. The chromosomal-length *D. citri* genome assembly serves as a blueprint for the development of collaborative genomics projects for other medically and agriculturally significant insect vectors.

## 1. Background

High throughput methods in genomics, transcriptomics and proteomics coupled with recent developments in single cell technologies have allowed for better understanding of processes at the cellular, organismal and ecological levels. However, a high quality genome assembly and annotation of both coding and non-coding genes is critical to the success of ‘omics assays, as well as interdiction methods that are based on gene targets. Easily accessible genomic and transcriptomic resources and web-based tools are essential for developing therapies and finding long-term management strategies, especially for difficult diseases of agriculture. Huanglongbing is an example of how genomic and transcriptomic resources [1] can help provide real management strategies for an otherwise terminal agricultural industry.

Huanglongbing (HLB), also known as citrus greening disease, is the most devastating of all citrus diseases and there is currently no effective control strategy. In the state of Florida alone, HLB has caused over $7 billion in lost revenue since 2005 and thousands of lost jobs [2]. HLB affects all genotypes of *Citrus* and some other Rutaceae species, causing early fruit drop, uneven ripening, unpleasant fruit taste, leaf mottling, and tree death in 3-5 years. In Asia, North and South America, Oceania and the Arabian Peninsula, HLB is caused by a phloem-limited, gram-negative bacterium, “*Candidatus* Liberibacter asiaticus” (*C*Las) and its insect vector, the Asian citrus psyllid, *Diaphorina citri* Kuwayama (Hemiptera: Liviidae) moves the pathogen from tree to tree. Asymptomatic but infectious trees and minimal vector control have contributed to the rapid spread of *C*Las, making disease monitoring and control difficult, emphasizing the need to find vector-control strategies to stop or slow the spread to new, healthy trees [3]. While the vector-pathogen interactions of HLB have been reviewed and compared in multiple articles [4, 5], this study aims to update the current genomic resources available for the vector, *D. citri*.

A short read-based genome assembly (Diaci v1.0) from a Florida *D. citri* population has been an important resource for the HLB research community since 2011, with multiple updates and improvements adding to it over the following 10 years (Diaci v1.1, Diaci v1.9, and Diaci v2.0) [6]. The early Diaci v1.1 assembly proved to be fragmented, with a contig N50 of 34.4 Kb and scaffold N50 of 109.8 Kb. Moreover, a benchmarking universal single-copy orthologs (BUSCO) [7] analysis indicated that a significant number of conserved single-copy markers were missing, deeming improvements necessary. In 2018, a higher quality genome assembly (Diaci v2.0) was released on citrusgreening.org [1, 8] using PacBio and Dovetail technologies, reducing the contig count and improving the contig N50 to 749.5 Kb. Ongoing improvements to the genome including community-driven manual annotations [9] and transcriptomic and proteomic analyses of psyllid mRNAs and proteins differentially expressed during *C*Las transmission have also identified a number of mis-assemblies such as tandem duplications and a high degree of gene fragmentation [10]. The Diaci v1.1 assembly was also lacking in key areas that are needed for a thorough understanding of *C*Las-*D. citri* interactions, as well as interactions between *D. citri* and its primary, obligate, and facultative intracellular endosymbionts.

*D. citri* harbors two obligate bacterial endosymbionts with highly reduced genomes (“*Candidatus* Profftella armatura” and “*Candidatus* Carsonella ruddii”) within a specialized organ called the bacteriome, which evolved specifically to host these symbionts. As shown by their “*Candidatus”* status, “*Ca.* P. armatura” and “*Ca.* C. ruddii” are unculturable and are limited to one specific organ within *D. citri*. The endosymbionts of *D. citri* are critical to psyllid survival and reproduction - “*Ca.* C. ruddii” produces essential vitamins allowing *D. citri* to feed from the nutrient poor phloem [11–14]. Suppression of “*Ca.* P. armatura” in *D. citri* resulted in increased psyllid mortality [15, 16] and the enosymbiont’s defensive properties [17] are also of interest for vector control strategies. *D. citri* is also host to a specific strain of the facultative endosymbiont, *Wolbachia* (*w*Di), though its role has yet to be clearly determined [18, 19]. Separating these endosymbionts from the insect host can only be done computationally, and typically the endosymbiont genomes must be removed from sequencing data prior to *de novo* assembly or other downstream analyses of the *D. citri* genome. Because psyllid biology is intimately connected to, and regulated by, the bacteriome symbionts, strategies to target and suppress psyllid endosymbionts are emerging as treatments that can reduce psyllid vector populations while limiting the damage to non-target species. The endosymbionts also interact with pathogens and can affect the acquisition and transmission of the bacterial pathogen *C*Las [15,16,20].

Here, we report an improved assembly of a Florida *D. citri* genome which we call Diaci v3.0. Diaci v3.0 has chromosomal-length scaffolds (produced by *de novo* assembly of PacBio long reads and Dovetail Chicago and Hi-C scaffolding) and a new Official Gene Set v3 (OGSv3, generated by automated gene prediction using Illumina RNAseq and PacBio Iso-Seq data as evidence). This genome assembly is most notable for the high-quality gene annotation effort conducted in parallel. Through a community annotation effort, primarily involving undergraduate students mentored by senior scientists, gene sets (including chitin metabolism, cuticle formation, segmentation and segmental identity, signal transduction, protein degradation, chromatin remodeling, phototransduction, circadian rhythm, carbohydrate metabolism, melanization and spermatogenesis) were manually curated using the Apollo annotation editor [21].

In order to better understand genome architecture, we compared Diaci v3.0 to that of the hackberry petiole gall psyllid, *Pachypsylla venusta,* and to three *D. citri* genomes that were reported after our genome was publicly released [22]. We also describe a comprehensive and genome-independent *de novo* transcriptome based on a variety of available transcript evidence from different tissues and conditions. Additionally, high quality and complete genome assemblies for the endosymbionts “*Ca.* P. armatura” and “*Ca.* C. ruddii”, and high quality draft genomes for two *w*Di strains, provide genomics resources to further understand complex organ-specific, intermicrobial interactions.

## Data Description

### Background and Purpose of data collection

High quality genome assemblies and accurate annotation of both coding and non-coding genes are critical to the success of ‘omics assays. While the vector-pathogen interactions of Huanglongbing (Citrus Greening disease) have been reviewed and compared in multiple articles [4, 5], this study aims to update the current genomic and transcriptomic resources available for the vector, *Diaphorina citri*.

### General methods

DNA was obtained from a *D. citri* colony originating from psyllids collected in Indian River County, Florida and maintained in the U.S. Horticultural Research Laboratory, USDA, Fort Pierce, Florida. DNA for genome assembly was extracted from pooled *D. citri* adults and sequenced using PacBio long read technology followed by multiple rounds of assembly and scaffolding, followed by duplication reduction, error correction and exclusion of non-target organism reads produced the Diaci v3.0 genome assembly. Genome assemblies of three *D. citri* endosymbionts were generated using reads excluded during the Diaci v3.0 assembly process, then cleaned and verified. We visually compared synteny of Diaci v3.0 with another psyllid genome, and with another recently published *D. citri* genome assembly. Our Iso-Sequencing used RNA sequencing data and an established computational pipeline to assemble gene isoforms independent of the genome. Additionally, a *de novo D. citri* transcriptome was generated using publicly available short read RNA sequencing data, and was combined with the Iso-seq isoforms. Protein-coding genes were predicted and annotated following the MAKER annotation pipeline and results were informed using Mikado and RNAseq datasets. We also predicted Transcription factors and their motifs using previously published pipelines and performed preliminary analysis of the effect of *C*Las infection on transcription factors and their targets. Our manual curation was a huge part of this genome project and involved teams of student annotators, graduate students and faculty across multiple institutions. We have previously published the annotation pipeline used [23]. Finally, we compared the predicted proteins of 12 different insect species to create orthogroups and assign GO terms to *D. citri* genes. See Figure 5 for an overview of the methods.

### Data Availability

The data sets supporting the results of this article are available in NCBI via BioProject accessions No. PRJNA29447 and PRJNA609978.

## 2. Analyses and Discussion

### 2.1. Genome assembly

The Diaci v.1.1 genome was generated and assembled with Illumina short reads, and thus had limitations to its usefulness for genomics-based research. Assembly of large eukaryotic genomes using short read technology results in fragmentation and misassembly of the genome because the short reads do not provide the contiguity required to assemble long and high-quality scaffolds [24]. The development of long read sequencing techniques, such as PacBio [25], and chromosome structure capture technologies such as Dovetail Chicago and Hi-C [26, 27], afforded an opportunity to generate a chromosomal-length *D. citri* genome to be used by researchers in the citrus greening community.

We generated 36.2 Gb of PacBio long reads. The Canu assembler was used to create a PacBio-only *de novo* assembly with 38,263 high confidence contigs, referred to as unitigs. The N50 of the unitigs (the length of the shortest contig for which longer and equal length contigs cover at least 50 % of the assembly) was 29 kb. Intermediate range scaffolding was performed with the Dovetail Chicago [24] method. Scaffolding with the HiRise assembler [24] based on paired-end reads from Chicago libraries with 22.53 X coverage of the *D. citri* genome increased the N50 from 29 Kb to 383 Kb. This step resulted in 12,369 joins and corrected 48 misassemblies in the unitig set. These Chicago scaffolds were subjected to gap-filling with PBjelly [28], available on GitHub [29] to create an intermediate Diaci v2.0 assembly with 1906 scaffolds and a scaffold N50 of 749.5 Kb (Table 1). A second round of long range scaffolding was performed on the non-gap-filled Chicago scaffolds using paired-end reads from a Hi-C library [30], which had higher coverage and larger inserts than the Chicago library. The HiRise assembler made 1003 joins in the Chicago assembly and increased the scaffold N50 from 383 Kb to 26.7 Mb.

**Table 1:** Diaphorina citri genome assembly statistics compared across three version releases. Comparison of statistics for three versions of the Florida Diaphorina citri (Diaci) genome: Diaci v1.1 [9], Diaci v2.0 (see Section 2.1) and the current version Diaci v3.0. BUSCO (Benchmarking Universal Single-Copy Orthologs) percentages were obtained by comparing each assembly to the Arthropoda (1013) gene set. Repeat percentages were calculated using RepeatMasker. *Each genome assembly is named as such: Diaci = <u>Dia</u>phorina <u>ci</u>tri; v = version. **13 chromosomal-length scaffolds with 1244 shorter non-redundant unplaced scaffolds, where shorter scaffolds are combined into a 14th “chromosome” with 1000 N’s between each scaffold. Additional, redundant and unplaced scaffolds which did not map to any of the current chromosomal-length scaffolds are not included in this number. ^Mb = Megabytes, Kb=Kilobytes.

| ***D. citri* genome assembly version\*** | **Diaci v1.1** | **Diaci v2.0** | **Diaci v3.0** |
| --- | --- | --- | --- |
| **Genome total size** | 485.7 Mb<sup>^</sup> | 498.8 Mb | 474 Mb |
| **Scaffold N50** | 109.8 Kb | 749.5 Kb | 40.6 Mb |
| **Maximum scaffold length** | 1.0 Mb | 4.2 Mb | 50.3 Mb |
| **Contig N50** | 34.4 Kb | 29.0 Kb | 29.0 Kb |
| **Maximum contig length** | 431.2 Kb | 2.1 Mb | 409.7 Kb |
| **Total scaffolds** | 161,988 | 1,906 | 1,257<sup>**</sup> |
| **Number of N's** | 19.3 Mb | 4.5 Mb | 13.4 Mb |
| **Complete BUSCO (arthropoda)** | 82.6 % | 92.1 % | 93.5 % |
| **Repeat %** | 26.4 % | 31.9 % | 30.2 % |

After Hi-C scaffolding, the assembly contained 13 chromosomal-length scaffolds (likely representing the 12 putative chromosomes [31] and the sex chromosome in the *D. citri* genome) and 24,930 unplaced scaffolds. We expected a high rate of duplication in the unplaced scaffolds in this assembly due to the pooling of multiple individuals to obtain the long read data, so we used Redundans [32] to classify the the unplaced scaffolds as unique or redundant compared to the rest of the genome assembly. We joined unique scaffolds (with 1000 Ns separating adjacent scaffolds) to create chromosome 00 in the Diaci v3.0 assembly. The 23,672 redundant scaffolds may represent alternative loci in the *D. citri* population and are reported as alternate (ALT) contigs. We subsequently removed 12 unplaced scaffolds from chromosome 00 and 2.3 Mb from chromosomal-length scaffold 9 that were identified as microbial contamination. The final Diaci v3.0 assembly has a total size of 474 Mb and an N50 of 40.6 Mb (Table 1). The complete history of *D. citri* genome sequencing can be found at citrusgreening.org [8].

#### Endosymbiont and mitochondrial assembly

The genomic DNA isolated from *D. citri* included bacterial sequences from which we obtained near complete assemblies for the obligate *D. citri* endosymbionts “*Candidatus* Profftella armatura” and “*Candidatus* Carsonella ruddii”, and for two strains of the facultative endosymbiont *Wolbachia*. We were able to identify specific regions of the *Wolbachia* genome that are shared between the two strains, as well as regions that were unique to each (Supplemental Figure 1). We compared our assemblies to those previously published [18, 33] through orthology analysis based on sequence identity and found the assemblies to contain the majority of orthogroups present in all published genomes (Table 2). We utilized the mitochondrial genome of *D. citri* sourced from China [34] to determine the mitochondrial sequences for our Florida *D. citri* samples.

**Table 2:**
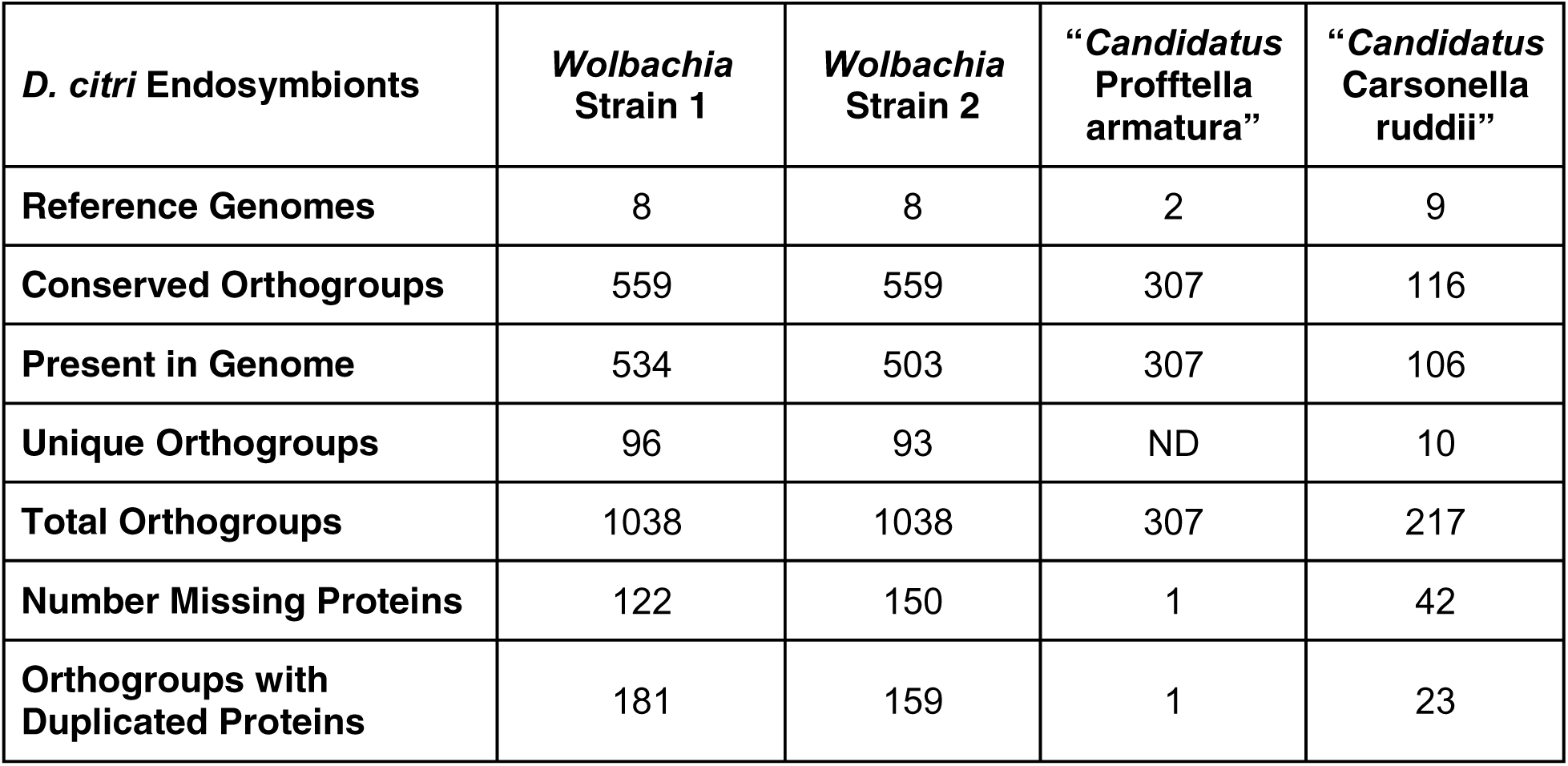
Genome statistics for *Diaphorina citri* endosymbionts including two obligate bacteria and two *Wolbachia* strains. An orthology analysis of endosymbiont genomes from Diaphorina citri compares the number of conserved orthogroups (the set of genes predicted to be derived from the last common ancestor) across published genomes to those present in our assemblies, as well as the number of orthogroups unique to our assemblies. We also present the number of proteins that may be duplicated or missing in each of our assemblies. Our assembly of the D. citri Wolbachia symbiont suggests two unique strains are present. The two other obligate symbionts are unculturable (thus, “Candidatus”) and play roles that are well documented in the literature. ND = no data.

### Comparative genomic analysis (synteny)

We compared the Diaci v3.0 assembly to the genome of the hackberry petiole gall psyllid, *Pachypsylla venusta,* [35] to identify regions of synteny (Figure 2). Three other members of the genus *Diaphorina* have a male 2N count of 25 (12 autosome pairs, plus the X chromosome) as is common among psyllids [36, 37]. The 13 chromosomal-length scaffolds in the *D. citri* genome assembly are consistent with these observations and match the number of chromosomal-length scaffolds in three recently published *D. citri* genomes [22]. In contrast, *P. venusta* has only 11 autosome pairs in addition to the X chromosome [38]. Most of the chromosomal-length scaffolds showed a one-to-one correspondence between the two species, suggesting their overall genome structure is highly conserved. The one major exception is that approximately half of *P. venusta* chromosome 3 (pv3, Figure 2) aligns with *D. citri* chromosomal-length scaffold 12, while the other half shares synteny with *D. citri* chromosomal-length scaffold 13 (dc12 and dc13, Figure 2). Thus, it seems likely that the reduction in autosome number in *P. venusta* is due to a fusion event between the chromosomes homologous to *D. citri* scaffolds 12 and 13.

**Figure 1.**
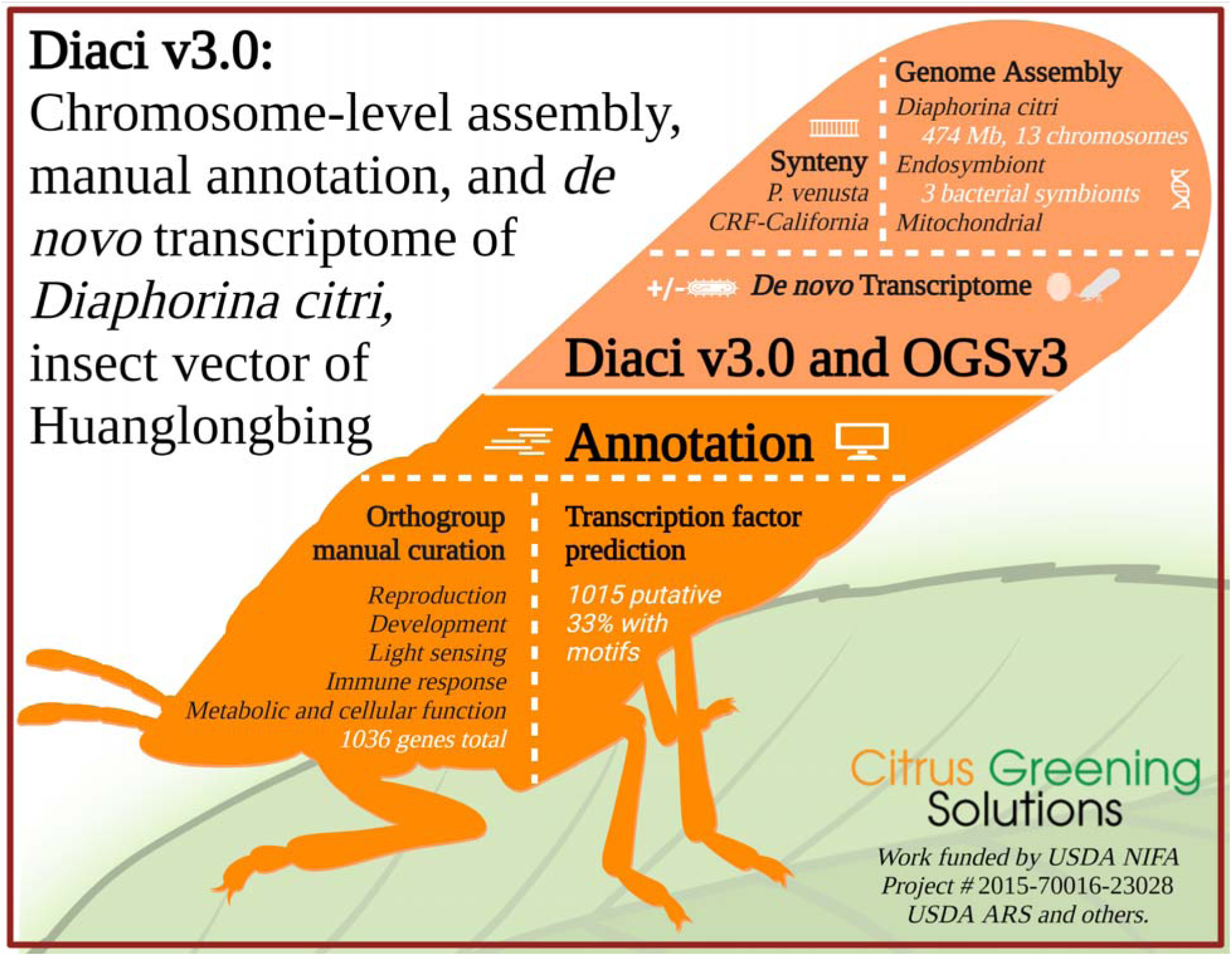
Graphical abstract of the major findings from a multi-year, multi-institutional *Diaphorina citri* genome assembly project. The most recent version (3.0) of the *Diaphorina citri* genome assembly (Diaci v3.0) is available on CitrusGreening.org and NCBI, along with its Official Gene Set v3, *de novo* transcriptome and extensive manual annotations covering major pathways and hundreds of genes. We have also predicted transcription factors and protein-coding genes, and compared Diaci v3.0 to the *Pachypsylla venusta* psyllid genome, as well as other *D. citri* genome assemblies recently published. Lastly, during our genome assembly, we also created draft genomes for multiple *D. citri* endosymbionts. Our work took place over eight years, multiple institutions and dozens of students, graduate students and faculty.

**Figure 2.**
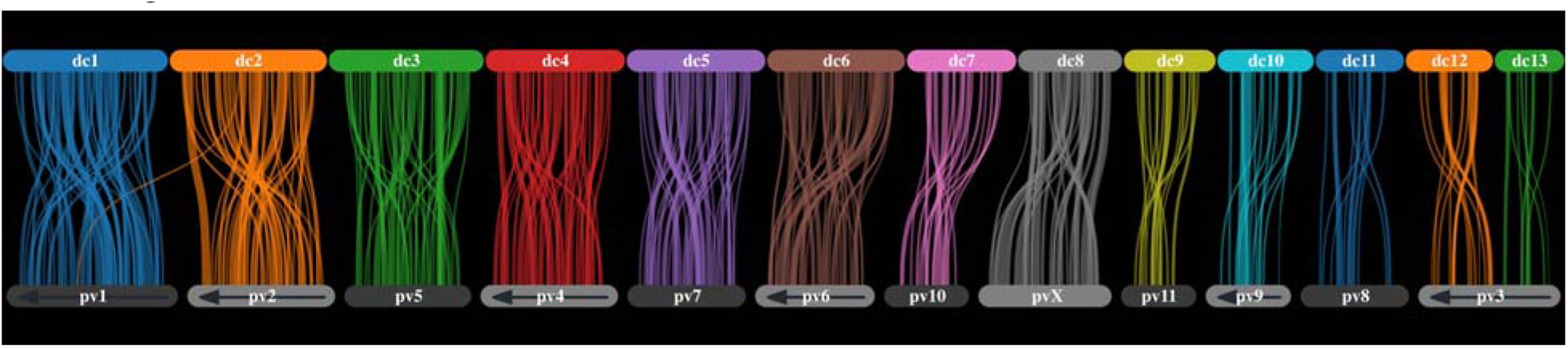
Parallel plot showing synteny between *Diaphorina citri* Diaci v3.0 and *Pachypsylla venusta* genome assemblies. *Diaphorina citri* chromosomal-length scaffolds from Diaci v3.0 are numbered in order of size (dc1-dc13) and make up the top row in the parallel plot. *Pachypsylla venusta* (the Hackberry petiole gall psyllid) chromosomal-length scaffolds are numbered as previously described (pv1-pv11) [35]. The *P. venusta* scaffolds marked with arrows (pv2, pv3, pv4, pv6, and pv9) have been reversed to optimize synteny to Diaci v3.0. We see that dc8 is likely the *D. citri* X chromosome, while pv3 appears to be a fusion of sequences homologous to dc12 and dc13.

We also used conserved synteny with *P. venusta* to identify the *D. citri* sex chromosome. The identity of the X chromosome scaffold of *P. venusta* has been established by comparing read mapping coverage of genomic sequence data from single males and females [35]. *D. citri* chromosomal-length scaffold 8 (dc8) is highly syntenic with the *P. venusta* X chromosome (pvX, Figure 2), allowing us to identify it as the probable *D. citri* X chromosome. Carlson et al [22] also compared one of their *D. citri* genomes with *P. venusta* and obtained the same results, identifying scaffold 7 of the CRF-California genome (which corresponds to scaffold 8 in the Diaci v3.0 genome, see Figure 3) as the *D. citri* X chromosome.

**Figure 3.**
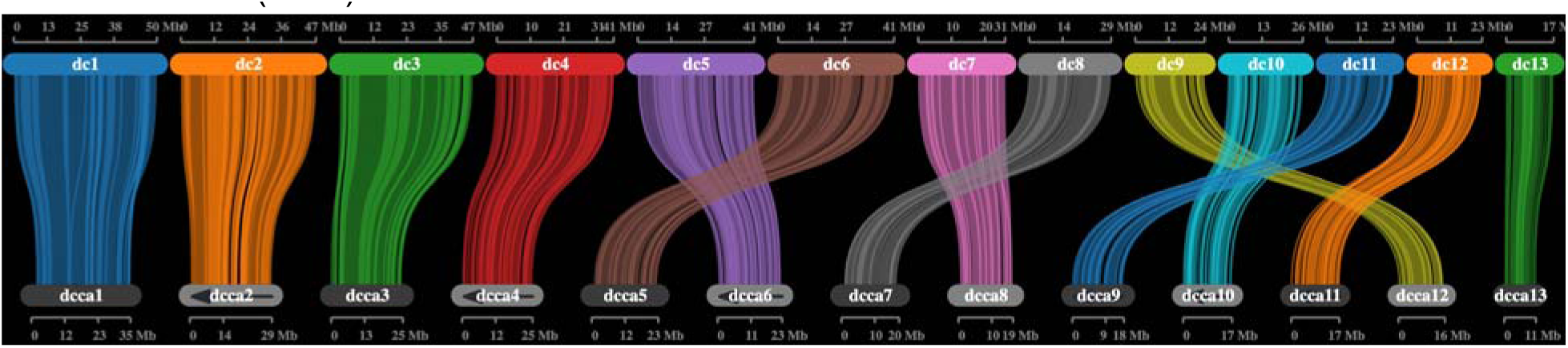
Parallel plot of two chromosomal-length *Diaphorina citri* genomes: Diaci v3.0 (dc) and CRF-California (dcca). The 474 Mb *Diaphorina citri* Diaci v3.0 Florida genome is about 200 Mb larger than the CRF-California (282.67 Mb) genome from Carlson et al [22]. All Diaci v3.0 chromosomal-length scaffolds (dc) are larger than the corresponding scaffolds of the CRF-California genome (dcca), suggesting that the additional sequence in the Diaci v3.0 assembly is distributed throughout the genome. Lines connecting scaffolds indicate syntenic blocks of genes. Scaffold sizes are shown in megabytes (Mb). Gray ovals with arrows indicate CRF-California scaffolds that have been reversed to match the orientation of the corresponding Diaci v3.0 scaffold.

Despite both efforts yielding chromosomal-length scaffolds, the 474 Mb Diaci v3.0 Florida genome is about 200 Mb larger than each of the three *D. citri* genomes described recently by Carlson et al [22]: CRF-Uruguay (266.76 Mb), CRF-California (282.67 Mb), and CRF-Taiwan (282.89 Mb). Moreover, flow cytometry analysis in the same paper gave genome size estimates of 274.4 Mb for males and 287.0 Mb for females. To investigate the reasons for the apparently inflated size of the Diaci v3.0 genome, we used MCScanX to align syntenic blocks between the Diaci v3.0 assembly and the CRF-California genome. The two genomes show one-to-one correspondence of chromosome-length scaffolds, with a few differences in numbering due to differences in relative scaffold size. All Diaci v3.0 scaffolds are larger than the corresponding scaffolds of the CRF-California genome, suggesting that the additional sequence in the Diaci v3.0 assembly is distributed throughout the genome (Figure 3). These results along with close examination of the v3.0 genome during manual annotation suggest that the Diaci v3.0 genome contains many heterologous loci that have been assembled into the main scaffolds as false tandem duplications. This issue likely stems from the use of multiple psyllids for DNA isolation during the Diaci v3.0 genome assembly process, a standard procedure at the time.

### 2.2. Annotation

We identified more repeats in the Diaci v3.0 genome assembly compared to the previously published Diaci v1.1 assembly (Table 1). This can largely be attributed to the use of long-read PacBio sequences in the Diaci v3.0 assembly. Computational annotation of genes was carried out with the MAKER annotation pipeline after masking repetitive elements. In total, we identified 18,947 protein-coding genes with 21,231 alternatively spliced isoforms (Table 3) in the Official Gene Set version 3 (OGSv3) gene annotation. OGSv3 includes automatically predicted as well as manually curated gene models. Use of PacBio Iso-Seq enabled accurate predictions of complete gene models and their isoforms. Predicted protein-coding genes were longer compared to OGSv1 gene models, with more exons predicted per gene (Table 3). Curated genes represent models that were manually refined using the Apollo annotation editor [21], based on evidence from orthology and expression data. Curation was performed by distributed groups based on the previously described workflow [23, 39]. Detailed reports of curated gene families in pathways of interest have been published and are summarized in section 2.5.2 below. Overall, curated genes are longer with more exons compared to predicted genes (Table 3). We also observe fewer non-canonical 5’ and 3’ splice sites in the curated genes and OGSv3 gene models, indicating higher quality of gene models compared to previous annotations. During the curation process, we removed 258 predicted genes that were not supported by expression data or represented false duplications. In addition to manual curation, we also generated functional annotation for OGSv3 gene models using the AgBase functional annotation pipeline [40] to assign experimentally validated GO terms, InterProScan domains and pathways based on sequence similarity.

**Table 3:** *Diaphorina citri* annotated gene counts for Official Gene Sets v1, v2, v3 and manually curated. The Diaphorina citri Official Gene Set versions 1, v2 and v3 include automatically predicted gene models (columns “OGSv1”, “OGSv2” and “OGSv3”), of which, over 1000 have been manually curated (column “Curated”). bp = base pairs. avg = average.

| Annotation Set | OGSv1 | OGSv2 | OGSv3 | Curated |
| --- | --- | --- | --- | --- |
| **Total Genes** | 19,311 | 20,793 | 18,947 | 1,036 |
| **Total Transcripts** | 20,966 | 25,292 | 21,231 | 1,143 |
| **Average Transcript Length (bp)** | 1,317 | 1,944 | 2,034 | 2,503 |
| **Exons Per Transcript (avg)** | 5.42 | 7.06 | 7.29 | 7.87 |
| **Average Exon Length (bp)** | 243 | 275 | 279 | 318 |
| **Non-Canonical Splice Sites** | 6.05 % | 3.13 % | 2.47 % | 1.91 % |

### 2.3. Transcription factor prediction

Transcription factors (TFs) are an important class of proteins that regulate gene expression levels by interacting with short DNA binding “motifs” in the genome. Knowledge of the TF repertoire and, when possible, the DNA binding motifs recognized by these TFs, thus represents a critical component of any genomics toolkit. We identified a total of 1,015 putative TFs in Diaci v3.0 (Figure 4, panel A). This value is similar to other insects [41–44]. e were able to infer motifs for 337 (33 %) *D. citri* TFs based on orthology (Supplemental File 1), mostly based on available DNA binding specificity data from *D. melanogaster* (228 TFs), but also from species as distant as human (77 TFs) and mouse (12 TFs). Many of the largest TF families have inferred motifs for a substantial proportion of their TFs, including Homeodomain (93 of 104, 89 %), bHLH (57 of 63, 90 %) and Forkhead box (24 of 31, 77 %). As expected from other insect systems [41–44], the largest gap in binding specificity knowledge is for C2H2 zinc fingers (only 39 of 377, ∼10 %), which evolve quickly by shuffling the regions encoding the many zinc finger DNA binding domains contained in their protein sequences..

**Figure 4.**
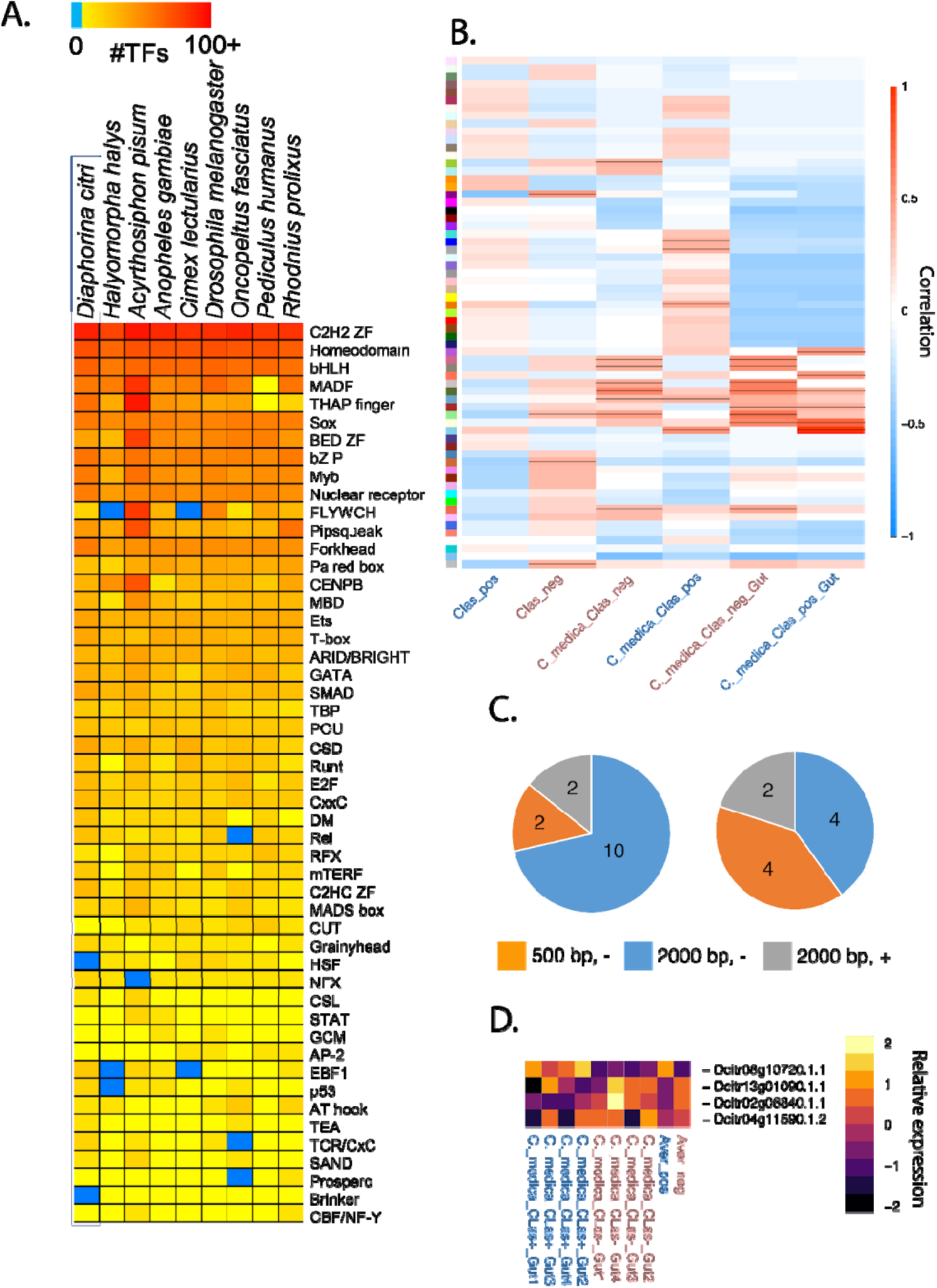
Comparison of transcription factors across insects and response to infection in *Diaphorina citri*. A) The top 50 families of transcription factors (TF, one shown per row) identified in *D. citri* (column 1) and in other insects (columns 2-9), where red is higher numbers of inferred TF motifs, and blue is no data. B) Weighted gene co-expression network analysis (based on FPKM values) and TF binding enrichment analysis identified specific modules of gene targets associated with *“Candidatus* Liberibacter asiaticus” (*C*Las) infection. Association between modules are shown as colors on the y-axis of co-expressed genes. CLas_pos are all *C*Las-positive (infected) and Clas_neg are all *C*Las-negative (healthy) samples. Samples denoted as C_medica_Clas_pos are CLas-infected *D. citri* raised on *Citrus medica* plants and those as C_medica_Clas_neg are healthy, non-infected raised on healthy *Citrus medica* plants. Lastly, C_medica_Clas_pos_Gut and C_medica_Clas_neg_Gut are the same as above but data comes only from guts of *D. citri* in both treatments. Red indicates higher correlation between module and *C*Las treatment with a black line indicating significant correlation at P < 0.05. C) Expression patterns of four TFs of interest show increased enrichment in binding sites for genes with differential expression during *C*Las infection. Increased expression shown as lighter yellow and decreased expression in dark blue. Specific genes listed include: Dcitr02g06840.1.1 (*protein mothers against dpp*, *Mad*); Dcitr08g10720.1.1 (ortholog of *ventral nervous system defective, vnd*); Dcitr13g01090.1.1 (transcription factor *Myb*); Dcitr04g11590.1.2 (*suppressor of hairless, Su(H)*). Aver_pos and Aver_neg are averages of all four *C*Las-positive gut samples and all four *C*Las-negative gut samples. The averages show altered TF binding prediction and differential expression within the gut. Transcript levels in A), B), and C) are reported from the Psyllid Expression Network [1] using Diaci v3.0 and the *de novo* transcriptome.

**Figure 5:**
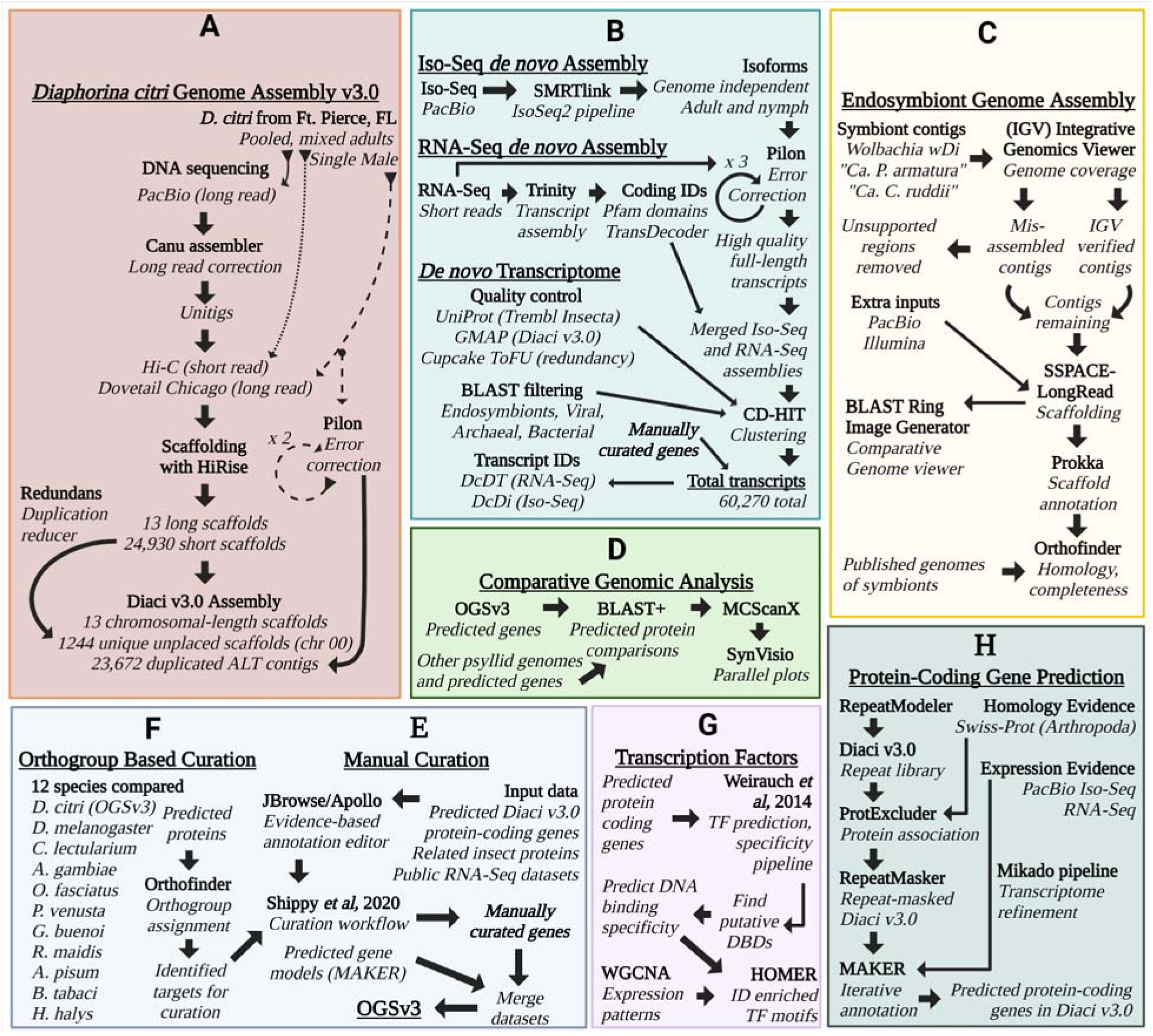
A visual representation of methods and pipelines for assembly, annotation, orthology, synteny, prediction and transcriptomics. A) The *Diaphorina citri* Diaci v3.0 genome assembly began with input of long read DNA sequencing data, followed by multiple rounds of assembly and scaffolding, duplication reduction, error correction and exclusion of non-target organism reads (see C). B) Iso-Sequencing used long read RNA sequencing data and an established computational pipeline (SMRTLink) to assemble gene isoforms independent of any genome. Additionally, a *de novo D. citri* transcriptome was generated using public short read RNA sequencing data collected from online repositories combined with the high quality Iso-seq isoforms. The transcriptome was assembled using Trinity, then cleaned and error corrected, subjected to BLAST to remove contaminants and clustered by locus to remove redundant isoforms. C) Genome assemblies of the *D. citri* endosymbionts including “*Candidatus* Profftella armatura”, “*Candidatus* Carsonella ruddii” and *Wolbachia* were generated using reads excluded during the Diaci v3.0 assembly process, then cleaned and verified using Orthofinder and BLAST. D) We performed synteny analysis between the Diaci v3.0 genome and other full length psyllid genome assemblies. E) Manual curation was a major part of this genome project and involved teams of student annotators, graduate students and faculty across multiple institutions. We have previously published the annotation pipeline used [23]. F) We compared the predicted proteins of 12 different insect species to generate orthogroups and assign GO terms to *D. citri* genes. G) Transcription factor (TF) prediction was performed following previously published pipelines and weighted gene co-expression network analysis (WGCNA) and HOMER were used to explore the effect of *C*Las infection on TFs and their target genes. H) Protein-coding genes were predicted and annotated following the MAKER annotation pipeline and results were informed using Mikado and RNAseq datasets.

Along with prediction of the TFs and their motifs, we used a combination of weighted gene co-expression network analysis (WGCNA) and TF binding enrichment analysis to identify specific gene targets of interest. The WGCNA identified specific modules of genes associated with infection with *C*Las (Figure 4, panel B). Based upon these modules, we used HOMER analysis [45] to identify the specific TFs whose binding sites were enriched upstream of the *C*Las-infection associated genes when compared to the whole gene set. From these analyses, there were 24 TFs with increased enrichment in binding sites for genes with differential expression in the whole body or the guts (Figure 4, panel C) within 2000 bp (Supplemental File 2) or 500 bp (Supplemental File 3) from the start of transcription. Transcriptional levels were found to be relatively high for four of these TF genes, with one increased and three decreased during *C*Las infection. Comparison with other insect systems revealed that the three genes with decreased expression are orthologs of *Myb*, *Mothers against dpp* (*Mad*), and *Suppressor of Hairless (Su(H))*, and the one that shows increased expression is the ortholog of *ventral nervous system defective (vnd)*. These TFs are involved in multiple aspects of development and could have critical roles in the response of *D. citri* to *C*Las and potential biological trade-offs that may occur during infection. As an example, *Drosophila melanogaster Mad* loss of function alleles result in defects in midgut and fat body morphogenesis [46], two organs known to be involved in *C*Las-*D. citri* interactions [14,47,48]. The other TFs identified have been associated with many processes that range from neuronal development to cell differentiation [49], but functional studies will be necessary to confirm the roles in relation to *C*Las-*D*. *citri*.

### 2.4 *De novo* Transcriptome

We identified a number of missing and incomplete genes during our detailed manual curation and orthology analysis of the *D. citri* genome, so we created a genome-independent *de novo* transcriptome to generate a more comprehensive set of transcripts. The transcriptome set was generated using 1.4 billion short and long individual reads from a range of experimental conditions and tissue types (Supplemental Table 3). This resulted in 40,637 genes and 60,261 transcripts with an average length of 1,736.1 bp and contig N50 of 3,657 bp. To differentiate transcripts in downstream analysis we assigned transcript identifiers according to the transcript evidence source. The DcDTr identifier prefix corresponds to 41,457 transcripts assembled from short read RNA sequencing data. And, the DcDTi identifier prefix was assigned to 18,804 transcripts supported by long read Iso-Seq data. Functional descriptions were assigned to 27,813 transcripts. We later added nine genome-independent transcripts (from seven genes) that were identified during manual curation. Most of these came from the MCOT transcriptome we described previously [6]. The transcriptome was validated with the hemipteran BUSCO data set and has 94.2% complete single-copy orthologs, and only 3.2 % missing single-copy orthologs (Supplemental Table 2).

### 2.5. Pathway-based manual curation

Community curation of *D. citri* genes was initiated with the Diaci v1.1 genome, resulting in more than 500 curated genes in Official Gene Set v1.0 [6], and has continued with subsequent genome versions. This initiative is particularly noteworthy because it is primarily driven by undergraduate students with input and supervision from scientists at multiple research institutions. The students were mentored by senior peer student annotators as well as expert annotators. Students had regular interactions with scientists from the insect genomics community working on genes and pathways under curation. Implementation of rigorous and consistent annotation practices across a virtual team of highly diverse annotators required project management tools and regular video conferences in addition to extensive documentation that was continuously updated in response to user feedback.

This group annotation strategy has been successful on multiple fronts (Table 4) and is likely to serve as a framework for integrating students into genomics to provide a high-quality genome resource for species of importance while educating the next generation of scientists in practical applications of bioinformatics methods. One of the most important lessons learned during this project is that our genome annotation and the student experience both benefit when students are recruited earlier in their academic career (first or second year students) and supported through a longer period than traditional undergraduate research experiences where students typically start research as juniors or seniors. These students can be involved from the start of analysis until publication, with a few generating their own first-author publications [50–55]. Students can join and contribute to genome annotation after first year introductory biology courses that cover basic molecular biology, which provides the necessary background to understand gene structure and function. The students can learn more complex biology through their annotation research experience with help from faculty and peer annotators. Students that are well trained and available to contribute for longer time frames reduce turnover and create a sense of a research community at local institutions that extends to having colleagues at other research institutions. Simply put, they learn the value of collaboration early in their careers and function with no silos around their mission and communication. Students that persist for two years or longer are extremely valuable for their ability to facilitate the training of new students through peer training and peer mentoring [6,39,56]. Several studies have shown that an extended duration for an undergraduate research experience increases student gains, both as part of the research project and individually in relation to general academic success [57]. This extended time frame also allows for higher level processing of data and the ability to interact with senior scientists, which helps students apply their knowledge of concepts and develop as independent research scientists [58]. Underrepresented students have been shown to specifically benefit from longer research experiences, with students reporting greater gains in scientific thinking, confidence to contribute to science, confidence working in teams and working independently [59].

**Table 4:** Overview of student genome annotation recruitment, training, and project outcomes. Across five institutions in multiple US states, dozens of undergraduate students, multiple graduate students, and a few faculty were involved in the manual curation of the predicted genes in the Official Gene Set version 3 (OGSv3). IRSC = Indian River State College (Florida), UC = University of Cincinnati (Ohio), KSU = Kansas State University (Kansas), Cornell = Cornell University (New York), BTI = Boyce Thompson Institute for Plant Research (New York), National level = US multi-state conference, Local/State = only one US state involved or a county-level local organization, STEM = Science, Technology, Engineering, Mathematics. ^Year of students: Typically undergraduate (Bachelors) students are enrolled two semesters a year, for four years (freshman, sophomore, junior, senior), while graduate students are post-Baccalaureate. *Publications with at least one undergraduate author include [6,23,50–55,60–62].

| **Institution** | **IRSC** | **UC** | **KSU** | **Cornell/BTI** |
| --- | --- | --- | --- | --- |
| Number of students | 53 | 32 | 11 | 4 |
| Year of students<sup>^</sup> | Second semester freshman to senior | Second semester freshman to senior | Freshman to senior | Freshman to senior |
| Number of Faculty/Senior scientists | 1 | 2 | 3 | 3 |

| Project delivery mode | Course,<br>Capstone,<br>Independent | Course,<br>Capstone,<br>Independent | Small<br>projects | Course,<br>Small<br>projects |
| --- | --- | --- | --- | --- |
| Annotation process | Pathways<br>and Gene<br>Families | Pathways<br>and Gene<br>Families | Pathways<br>and Gene<br>Families | Pathways<br>and Gene<br>Families |
| # Presentations (National level) | 1 |  |  |  |
| Presentations (local/state level) | 5 |  |  |  |
| Posters (national level) | 10 |  |  |  |
| Posters (local/state level) | 44 |  |  |  |
| Publications* | 12 |  |  |  |
| Careers (STEM professions) | 19 |  |  |  |
| Careers (Graduate/professional<br>school) | 18 |  |  |  |
| Total trainees (undergraduate) | 85 |  |  |  |
| Total trainees (graduate) | 3 |  |  |  |

#### 2.5.1. Orthogroup-based curation

In order to guide the manual curation, orthology analysis was performed with genes from 12 species (Bemisia tabaci, Acyrthosiphon pisum, Rhopalosiphum maidis, Diaphorina citri, Pachypsylla venusta, Drosophila melanogaster, Anopheles gambiae, Homalodisca vitripennis, Gerris buenoi, Cimex lectularius, Halyomorpha halys, Oncopeltus fasciatus) with 82.7 % of those genes being assigned to a total of 14,346 orthogroups. We utilized the orthogroups to confirm if a gene was missing from the Diaci v3.0 genome and verify if all the members of a gene family had been curated. The orthogroups also helped us identify duplicate (partial or complete) models that needed to be removed from the Official Gene Set v3 (OGSv3). We would like to note that using orthogroups is an effective strategy to initiate gene curation projects where an orthogroup containing a gene from Drosophila or another well characterized genome is selected followed by curation of the D. citri models in that orthogroup. Recently acquired genes from horizontal gene transfer or genes under positive selection that do not share sufficient sequence homology with orthologs will remain unclustered, but also offer valuable targets for manual curation.

#### 2.5.2. Specific pathways targets for curation

The community curation team, comprising undergraduate and graduate students and their mentors (Table 4 above), manually annotated several new sets of genes in the Diaci v3.0 official gene set, OGSv3, focusing our efforts on orthogroups, pathways of interest to the *D. citri* research community, and genes that have been targeted for RNAi-based pest control in other insects [63–65]. Detailed reports for these gene sets are included in 11 accompanying articles published as data releases by Gigabyte [50–55,61,62,66–68]. Sequences of the manually curated transcripts and predicted proteins are available for download at citrusgreening.org [8].

##### 2.5.2.1. Development

We annotated *D. citri* orthologs of many genes important for insect development, including genes involved in segmentation, signal transduction and segmental identity.

###### Segmentation

During embryonic development, the insect embryo is divided into segments. The process of segmentation differs somewhat between insects, but the genes involved are well conserved [69]. Out of 33 genes involved in *Drosophila* segmentation, we identified and annotated 25 in *D. citri* [67]. Most of the differences in gene content between *Drosophila* and *D. citri* were expected based on observations in other hemipterans [70, 71].

The segmentation genes are traditionally grouped into categories based on their function in *Drosophila*. In *D. citri*, we identified all of the maternal effect genes typically found in hemipterans, including *caudal*, *nanos*, *dorsal*, and one *TGFalpha*. As expected, we did not find *bicoid* or *oskar* [72]. We were unable to find several gap genes in *D. citri*, including *giant*, *buttonhead, orthodenticle 1* and *eagle*. All of these genes, except *eagle*, are also missing in the pea aphid genome. We did, however, find an ortholog of *huckebein*, which was reported absent in the pea aphid [70]. We identified one-to-one orthologs of each of the pair-rule genes in *D. citri*. The segment polarity genes include members of the Wnt and Hedgehog signal transduction pathways, which are described in the next section, as well as the transcription factor genes *engrailed* (*en*) and *gooseberry* (*gsb*). We identified a single copy of the *en* gene in *D. citri*, but we were unable to find its paralog *invected* (*inv*) despite the presence of both *en* and *inv* in other hemipterans [41,70,73].

###### Wnt and other signaling pathways

The Wnt signaling pathway is critical for segment polarity determination during segmentation and plays many other roles during development and in other biological processes [74, 75]. Particular ligand genes have been lost in various insects [74] and *D. citri* is no exception. We were unable to find orthologs of *Wnt8/D* or *Wnt9* [54], which are also absent in pea aphids. We also did not find *Wnt16*, although it is present in several other hemipterans [70, 76]. Conversely, *Wnt6* (contrary to a previous report [77]) and *Wnt10* are present in the *D. citri* genome but apparently absent in pea aphid [70]. The key downstream components of Wnt signaling are all present in the *D. citri* genome. We also annotated 13 genes from the Hedgehog pathway, 16 from the Notch pathway, and five members of the insulin signaling pathway. All of these pathways seem to be highly conserved in *D. citri*.

###### Hox genes and their cofactors

Hox genes specify the identity of regions along the body axis. Notably, they are usually arranged in a chromosomal cluster in an order paralleling that of their functional domains along the anterior-posterior body axis, although breaks in the cluster have occurred in several lineages [78]. We identified the full complement of ten Hox genes in *D. citri*, although *labial*, which was present in previous versions of the genome, is missing in Diaci v3.0 due to a local misassembly [66]. The *D. citri* Hox cluster is split into two parts with the breakpoint between *Dfd* and *Scr*, the first time a break between these genes has been reported in insects. Both clusters of Hox genes are on the same chromosome about 6 Mb apart.

We also annotated four TALE-class homeobox genes encoding proteins that frequently serve as Hox cofactors in other organisms [79]. *D. citri* has two members of the MEINOX class, which is consistent with other insects. However, while most insects have only one copy of the PBC class gene *extradenticle* (*exd*), *D. citri* has two. One of these *exd* genes has no introns and appears to be a retrogene. In *D. citri,* both *exd* genes are expressed in a wide range of stages and tissues, suggesting they could both be functional. Somewhat surprisingly, comparison to other insect Exd proteins shows that the protein encoded by the retrogene is more conserved than the protein encoded by the *exd* copy with a more typical gene structure, raising the possibility that the retrogene has taken over the function of the original gene. Functional studies of both genes could help determine which of the genes has retained the original function of *exd* and whether either has acquired any new functions.

##### 2.5.2.2. Immune Response

###### Melanization

The melanization pathway is important for wound healing and defense against pathogens, in addition to its role in pigmentation [80]. We annotated 12 genes from the melanization pathway including two laccases and two tyrosinase prophenoloxidases. We also annotated the members of the *yellow* gene family, several of which have been implicated in melanization in other insects, although their precise role is still not well understood [81]. *D. citri* has nine *yellow* genes, including an apparent duplication of *yellow-y* [55]. Interestingly, one of these paralogs is expressed primarily in the egg and nymph stages, while the other is expressed mainly in adults. We also noted apparent differences in expression of some *yellow* genes between *C*Las-infected and uninfected psyllids, warranting further investigation into their potential role in *C*Las transmission.

##### 2.5.2.3. Metabolic and Cellular Functions

Housekeeping genes involved in metabolic pathways and essential cellular functions are excellent targets for RNAi-based pest control, since knockdown is often lethal. We have annotated genes from several essential pathways including carbohydrate metabolism, chitin metabolism, chromatin remodeling, protein degradation and organelle acidification.

###### Carbohydrate Metabolism

Although trehalose is the primary blood sugar in insects, glucose metabolism is also important [82]. Breakdown of trehalose produces glucose, which is then further metabolized by glycolysis [83]. The synthesis of glucose and trehalose (gluconeogenesis and trehaloneogenesis, respectively) follow the same pathway up to the production of glucose-6-phosphate, before diverging. Some insects synthesize both sugars, with glucose synthesis occurring primarily in neural cells [82]. We annotated 32 genes involved in carbohydrate metabolism [52]. The genes in these pathways are highly conserved, although there are a few differences in gene copy number. Most notably, *D. citri* has two copies each of *phosphoenolpyruvate carboxykinase* and *fructose bisphosphate-aldolase* instead of a single copy as found in many other insects. We did not find a *glucose-6-phosphatase* gene in *D. citri*, suggesting that trehalose is the end product of this pathway in *D. citri*.

###### Chitin Metabolism

Chitin is a major component of the insect cuticle and properly coordinated synthesis and breakdown of chitin are essential for growth and survival [84]. We annotated 19 orthologs of genes involved in chitin metabolism: three genes for enzymes involved in chitin synthesis [68], four chitin deacetylase genes [61] and 12 genes whose products are involved in chitin degradation [62]. Like most hemipterans and other hemimetabolous insects, *D. citri* has fewer chitin metabolism genes than do most holometabolous insects. This reduction is likely due to the fact that hemipterans do not undergo complete metamorphosis and their guts lack a true peritrophic membrane (another structure containing chitin) [85]. Consistent with the absence of a peritrophic membrane, *D. citri*, like other hemipterans, lacks the chitin synthase 2 gene (CHS2), which is specifically expressed in the peritrophic membrane in holometabolous insects [86, 87]. *D. citri* and *A. pisum* [88] both lack a group VII chitinase, although one is present in some hemipteran species [89, 90]. *D. citri* has apparent duplications of two genes involved in chitin metabolism: UDP-N-acetylglucosamine pyrophosphorylase (UAP) and Chitinase 10 (a group II chitinase). We also identified a chitinase gene that seems to have arisen by horizontal gene transfer [62].

###### V-ATPase

Vacuolar ATP synthase (V-ATPase) regulates the acidity of various organelles by using energy from ATP to translocate protons across a membrane [91]. We annotated 14 genes encoding subunits of V-ATPase [53]. Gene copy number for each subunit varies between insects. The ACP genome has two copies of the V_0_-α subunit gene and one copy of each of the others. We also annotated one gene encoding an accessory V-ATPase subunit that may help assemble the V-ATPase complex.

###### Ubiquitin-proteasome pathway

Ubiquitination is an ATP-dependent process that targets proteins for degradation. We annotated 15 genes of the ubiquitin-proteasome pathway in the ACP genome [51]. These genes were chosen based on studies by [15], who identified several *D. citri* ubiquitin-proteasome pathway proteins as being differentially expressed during *C*Las infection, and [64], who found multiple ubiquitin-proteasome pathway genes among high lethality targets in a *Tribolium castaneum* RNAi screen.

###### Chromatin Remodeling

Chromatin remodeling proteins modify the positioning of nucleosomes along the chromosome to control the accessibility of particular regions of genomic DNA to protein binding. This is particularly important for regulating the transition between transcriptionally active and inactive chromatin states. We annotated 27 *D. citri* genes encoding chromatin remodeling proteins. Overall, the gene content of this family closely matches that of *Drosophila*. However, there is a duplication of *Mi-2* that has not been reported in other insects.

##### 2.5.2.4. Light sensing

Light sensing is essential for vision and maintenance of day-night rhythms. Both processes have a strong effect on the behavior of insects. To gain insight into these pathways in the psyllid, we searched for genes known to be involved in circadian rhythm and phototransduction.

###### Circadian Rhythm

We annotated 27 orthologs of genes putatively involved in circadian rhythm in other insects [50]. All components of the pathway are present in the *D. citri* genome. Notably, *D. citri* has two cryptochrome photoreceptor genes (*cry1* and *cry2*), which are believed to be the ancestral state of this pathway in insects [92–94].

###### Phototransduction

We annotated 21 genes in the phototransduction pathway. Opsin genes are important for the ability of an organism to detect light [95]. *D. citri* has four copies of opsin, the same number as the honeybee but fewer than the seven found in *Drosophila* [96]. Single gene copies of *UV-sensitive opsin, short-wavelength opsin, long-wavelength opsin*, and *rhodopsin 7* were annotated.

##### 2.5.2.5. Reproduction

Genes involved in reproduction, particularly spermatogenesis, have been targeted for use in a method of pest control called Sterile Insect Technique (SIT). Male insects sterilized by radiation, chemicals, or in this case, RNAi-based knockdown of spermatogenesis genes are released into the environment with the hope of out-competing fertile males in the natural population [97, 98]. We have annotated 20 *D. citri* orthologs of genes that have been identified as potential SIT targets in other insects [99–101].

### 2.6. Improving OGSv1 curated gene models

A subset of genes that had been manually annotated in the Diaci v1.1 genome [6] were examined in the improved Diaci v3.0 genome. Of 159 previously annotated genes, 27 were removed from the Official Gene Set (OGS) version 3 (Table 5). All but one of these were artifactual duplicates resulting from misassemblies in Diaci v1.1. Most of these duplications are no longer present in Diaci v3.0 due to long-read based assembly and removal of duplicate scaffolds as previously described. One heat shock gene was removed because it was located in a region of Diaci v1.1 determined to be of endosymbiont origin and was shown by BLAST to be identical to the molecular chaperone HtpG from “*Ca.* P. armatura”. Of 24 annotated gene pathways or families examined, eight have a reduced and more accurate gene count after validation in Diaci v3.0 (Table 5).

**Table 5.** Comparing Diaci v1.1 and Diaci v3.0 gene counts by pathway or gene family. Gene counts of manually curated gene families or pathways in Diaci v1.1 versus Diaci v3.0. The asterisk (*) indicates a case where one lysozyme gene was missing from Diaci v3.0 but was found in the ALT contigs.

| **Diaci v1.1 Pathway / Gene Family** | **Number of genes in Diaci v1.1** | **Number of genes in Diaci v3.0** |
| --- | --- | --- |
| Toll Receptors | 5 | 5 |
| Toll Receptor related genes | 3 | 3 |
| JAK/STAT pathway | 3 | 3 |
| C-type Lectins | 10 | 7 |
| Lysozymes | 5 | 5* |
| Superoxide dismutases | 4 | 4 |
| CLIP | 11 | 10 |
| Autophagy | 15 | 15 |
| Dicer | 4 | 2 |
| Drosha | 4 | 1 |
| Pasha | 2 | 2 |
| Loquacious and R2D2 proteins | 3 | 3 |
| Argonaute and PIWI proteins | 4 | 4 |
| Tudor Staphylococcal Nuclease protein | 2 | 1 |
| Vasa intronic gene | 1 | 1 |
| Armitage | 1 | 1 |
| Fragile X Mental Retardation Protein | 1 | 1 |
| Spindle_E (homeless) | 2 | 1 |
| Rm62 protein | 1 | 1 |
| Ras-related nuclear protein | 1 | 1 |
| Aquaporin | 6 | 6 |
| Cathepsins and Cysteine Proteases | 34 | 23 |
| Heat shock proteins | 18 | 13 |
| Rab genes | 19 | 19 |
| **Total** | **159** | **132** |

We were able to improve almost all re-examined models. In many cases, the old models had only partial open reading frames (ORFs) and we were able to create models with complete ORFs. We were also able to add 5’ and 3’ UTRs to many models. The Iso-Seq transcripts were extremely helpful in creating longer models with a high level of confidence.

We were also able to improve the accuracy of annotated transcriptional isoforms in Diaci v3.0. We discarded 12 models previously annotated as differentially spliced isoforms. In most cases, these errors stemmed from falsely duplicated exons in the Diaci v1.1 genome being interpreted as alternative exons. Although our efforts were focused on assessing existing models, we did identify and annotate two previously unrecognized isoforms in the Diaci v3.0 genome during the validation process.

## 3. Conclusions and Potential Implications

The hemipteran *Diaphorina citri* (Asian citrus psyllid) is a primary target of approaches to stop the spread of the bacterial pathogen *C*Las that causes Huanglongbing (citrus greening disease) (reviewed in the book by Qureshi and Stansly (2020) [102], Chapters 11-17). In support of the global effort to control this insect vector, we provide a significantly improved, chromosomal-length genome assembly of the *D. citri* genome (Diaci v3.0), along with the corresponding Official Gene Set (OGSv3) which includes 18,947 protein coding genes, of which 1036 have been manually curated. The hologenome of this species is provided by direct sequencing of the two microbial symbionts “*Ca.* Profftella armatura” and “*Ca.* Carsonella armatura”, and the *D. citri* strain of *Wolbachia* (wDi).

Under this project, we have also advanced the education and expertise of undergraduate and graduate students from multiple institutions who received training in gene curation as part of the community curation effort. Many trainees were also involved in the production of concrete deliverables in the form of research publications and presentations, including previous genome releases, this paper, and other direct studies completed as part of this project [10,50–55,61,62,66–68]. Standard operating procedures have been reported as part of a guide for other annotation communities [23, 39] to implement similar programs.

All resources are available online at CitrusGreening.org, a portal for ‘omics resources for the citrus greening disease research community [1]. The chromosomal-level *D. citri* genome assembly, automated annotation based on transcriptomics evidence, manual curation of critical pathways and a genome-independent *de novo* transcriptome will provide a foundation for comparative analysis among genomes of agricultural pests and plant-feeding hemipteran vectors of plant pathogens. Moreover, these resources will facilitate genome-wide association studies to identify the *D. citri* genes involved in *C*Las acquisition and transmission.

## Methods

### 1. *Diaphorina citri* Genome assembly

The DNA for PacBio sequencing and the Dovetail Chicago and Hi-C libraries was sourced from a *D. citri* colony originating from psyllids collected in Indian River County, Florida and maintained at the U.S. Horticultural Research Laboratory, USDA, Fort Pierce, Florida. High molecular weight DNA for PacBio sequencing was extracted from pooled *D. citri* adults using previously described methods [103]. DNA from a single male *D. citri* was used to generate the Dovetail Chicago library which was sequenced and used for both scaffolding and error correction.

PacBio sequencing was done on an RSII instrument to produce 36.2 Gb of long reads with an average length of 7.2 Kb. The Canu [104] assembler software was used to correct 40 X of the longest continuous long reads which were trimmed and used for the final assembly (-utgOvlErrorRate=0.013). The 38,263 unitigs produced by the Canu assembly with a contig N50 of 29 Kb were selected for scaffolding. 207 million Dovetail Chicago paired-end reads from a single male psyllid, with an insert size distribution of up to 250 Kb, were used to perform 12,369 joins and connect these unitigs into 25,942 scaffolds. This round of scaffolding added 12.3 Mb of Ns to the assembly. The Chicago scaffolded assembly was passed through another round of scaffolding with 388 million paired-end Hi-C reads [105]. The insert size distribution of the paired-end reads from the Hi-C library stretched up to 3 Mb with genome coverage of 819.14 X. The Hi-C scaffolding and validation reduced the number of scaffolds to 24,943 containing 12.4 Mb of Ns and a scaffold N50 of 26.7 Mb. This assembly consisted of 13 chromosomal-length scaffolds comprising 441 Mb of the genome. There were also 24,930 “unplaced scaffolds” that were not placed into the putative chromosomes. This unplaced set consisted of short scaffolds (scaffold N50 15 Kb). We reduced the duplication within the unplaced sequences by applying Redundans [32] at a threshold of 80 % identity and coverage. This split the unplaced set into 1271 unique scaffolds (33.2 Mb, N50 28.1 Kb) and 23,672 duplicated scaffolds (201 Mb, scaffold N50 13.7 Kb). The duplicated scaffolds are reported as alternate (ALT) contigs for the Diaci v3.0 assembly. Unique scaffolds were ordered based on length and joined with 1000 Ns separating adjacent scaffolds to create chromosome 00 of length 34.5 Mb. We performed two rounds of error corrections with Pilon [106] using Illumina reads from the single male psyllid individual. We opted to use this data set instead of short read data from multiple individuals to avoid introducing artificial heterozygosity into the genome assembly. Pilon was optimized to only correct regions of the genome where the change was supported by more than 90 % of the aligned bases at that position (--fix bases --diploid --mindepth 0.9). We also performed one round of error correction with Pilon and Illumina RNA-Seq to polish the genic regions of the assembly (--unpaired --fix bases --diploid) based on unspliced alignments to the genome. To minimize microbial contamination, we removed regions of the assembly with very high read coverage (>1000x) and BLAST hits (>200 bp and >90% coverage) to an endosymbiont genome. We used BUSCO [7] Arthropoda (arthropoda_odb10 with 1013 markers) and Hemiptera (hemiptera_odb10 with 2510 markers) marker sets to evaluate the completeness of all the data sets.

#### 1.1. Endosymbiont assembly, validation and annotation

To identify endosymbionts present in *D. citri*, we identified contigs from the Diaci v3.0 assembly with sequence similarity to published “*Ca.* P. armatura”, “*Ca.* C. ruddii”, and *w*Di genomes. Endosymbiont contigs were manually validated using the Integrative Genomics Viewer (IGV) [107] to assess coverage of PacBio and Illumina DNA reads. Any mis-assembled contigs were split into multiple contigs and regions that had no support were removed from further analysis. We applied the SSPACE-LongRead [108] scaffolding algorithm to contigs verified in IGV in order to join any contigs for which there was PacBio evidence of overlap. When running SSPACE we required 20 PacBio reads to link contig-pairs for scaffolding (-l 20). We annotated scaffolds with Prokka [109] and analyzed them using Orthofinder [110] and BLAST Ring Image Generator (BRIG) [110, 111]. We used Orthofinder to judge the completeness of the genome assemblies by comparing the orthogroups present in our assemblies to those present in published genomes. We defined a conserved orthogroup for a given endosymbiont to be one that contains at least one protein from each reference genome (Supplemental Figure 1). A conserved orthogroup then indicates protein functionality shared among all known genomes. We defined a unique orthogroup for a given endosymbiont to be one that is present in our assembly, and is not present in at least 50 % of the reference genomes.

#### 1.2. Comparative genomic analysis

We used MCScanX [112] to identify syntenic blocks between the chromosomal-length scaffolds of the Diaci v3.0 genome and other publicly available psyllid genomes [22, 35]. Simplified GFF files of predicted genes from each genome were prepared by using gffread v0.12.1 [113] to extract scaffold names, gene names and coordinates from available GFF files. For each genome pair, BLAST+ [114] was used to compare predicted proteins in all possible pairwise combinations. The BLAST results and simplified GFF files were then used as input for MCScanX with default parameters. Parallel plots were created from the MCScanX results using SynVisio [115].

### 2. Iso-Seq

In order to detect alternative splicing we performed isoform sequencing (Iso-Seq) using PacBio to generate high quality (HQ), full length transcripts. Samples were collected from healthy and *C*Las-infected adult *D. citri*, and healthy and *C*Las-infected nymph *D. citri*. The sequences were processed using the PacBio SMRTlink v4.0 software [116]. The pipeline runs circular consensus sequencing (CCS) using the raw reads followed by a classification of full length and non-full length transcripts. The resulting transcripts were polished to generate HQ consensus isoforms and polished low quality (LQ) consensus isoforms. A comprehensive set of genome independent HQ Iso-seq isoforms from adult and nymph tissue was achieved with three rounds of error corrections with Pilon [106] using Illumina RNA-Seq reads.

### 3. *De novo* transcriptome

The *de novo* transcriptome was generated using the Iso-Seq data described above and publicly available short read RNA-Seq data from different experimental conditions and tissues, obtained from the NCBI SRA database (Supplemental Table 1).

Trinity [117] was used to perform *de novo* assembly of both RNAseq and Iso-Seq datasets using default parameters. Transcripts that did not encode at least one Pfam domain [118] or did not contain a coding region (as determined by TransDecoder [119]) were removed from the dataset. Sequences were then clustered with CD-HIT (cd-hit-est) [120] using a threshold of 75% sequence identity, and the longest transcript in each cluster was selected. We used BLAST [121] to identify contamination from archaeal, viral and bacterial sequences and removed those transcripts. We also removed transcripts that did not have any matches to the Trembl Insecta subset in the Uniprot database [122]. Finally, the remaining transcripts were mapped to Diaci v3.0 using GMAP [123] and the Cupcake ToFU [124] pipeline was used to collapse redundant isoforms at each locus. We assigned transcript identifiers according to the source of the raw data: DcDTr for RNA-Seq transcripts and DcDTi for Iso-Seq transcripts.

### 4. Automated predictions of protein-coding genes

A repeat library for the Diaci v3.0 genome was constructed using RepeatModeler [125]. The repeat library was screened for known protein association with ProtExcluder [126] based on similarity with proteins obtained from Swiss-Prot (Arthropoda) [122]. The resulting repeat library was used to mask the Diaci v3.0 genome with RepeatMasker [125]. Repeat annotation is available on citrusgreening.org FTP.

Protein-coding genes were predicted on the repeat-masked Diaci v3.0 genome through iterative processing within the MAKER (v3) annotation pipeline [127]. For homology evidence, manually annotated proteins of Arthropoda were downloaded from Swiss-Prot [122]. Expression evidence was obtained through multiple sources. RNA-Seq data generated as part of this work is listed in Supplemental Table 1. Publicly available RNA-Seq datasets were obtained from NCBI SRA database (Supplemental Table 1). All the RNA-Seq data was mapped to the genome using HISAT2 [128] and the transcriptome was assembled with StringTie [129]. Independently, high-quality PacBio Iso-Seq transcripts were mapped to the genome and clustered through Cupcake-ToFU clustering [124]. Transcriptomes obtained through RNA-Seq and Iso-Seq were processed with Mikado pipeline [130] for refining the transcriptome.

*Ab initio* gene predictions were performed with Augustus [131] and SNAP [132] gene predictors. Augustus was trained with available RNA-Seq data within BRAKER1 [133]. SNAP was trained iteratively within the MAKER pipeline based on MAKER guidelines. The gene predictions were supplied to MAKER along with expression and homology evidence which was run with default parameters. The Mikado-refined transcriptome was passed as a predictor (pred_gff). Gene identifiers were assigned based on their genomic location. For example, the Dcitr01g01000.1.1 gene consists of a five letter species identifier (Dcitr), a two digit scaffold/chromosome number (01), an abbreviation for gene (g), a unique five digit ID for the gene (01000) followed by version number (.1) and isoform number (.1). Consecutive genes were assigned identifiers in increments of 10 to allow addition of genes in the future.

### 5. Transcription factor prediction

We identified likely transcription factors (TFs) by scanning the amino acid sequences of predicted protein coding genes for putative DNA binding domains (DBDs), and when possible, we predicted the DNA binding specificity of each TF using the procedures described in Weirauch *et al.* [134]. Briefly, we scanned all protein sequences for putative DBDs using the 81 Pfam [118] models listed in Weirauch and Hughes [135] and the HMMER tool [136], with the recommended detection thresholds of per-sequence Eval < 0.01 and per-domain conditional Eval < 0.01. Each protein was classified into a family based on its DBDs and their order in the protein sequence (e.g., bZIPx1, AP2×2, Homeodomain+Pou). We then aligned the resulting DBD sequences within each family using ClustalOmega [137], with default settings. For protein pairs with multiple DBDs, each DBD was aligned separately. From these alignments, we calculated the sequence identity of all DBD sequence pairs (i.e. the percent of AA residues that are exactly the same across all positions in the alignment). Using previously established sequence identify thresholds for each family [134], we mapped the predicted DNA binding specificities by simple transfer. For example, the DBD of Dcitr04g16960.1.1 is 98 % identical to the *Drosophila melanogaster* ‘oc’ (FBgn0004102) TF. Since the DNA binding specificity of ‘oc’ has already been experimentally determined, and the cutoff for the Homeodomain family of TFs is 70 %, we can infer that Dcitr04g16960.1.1 will have the same binding specificity as ‘oc’ (Supplementary File 1). WGCNA [138] was conducted according to methods previously adapted to examine gene expression patterns in relation to TF binding predictions [44,139,140]. Enriched TF binding motifs were identified in the 500 and 2000 bp regions upstream of the putative transcription start site using the HOMER tool as in previous studies examining the dynamics between TFs and expression patterns in insects [45].

### 6. Manual curation and OGSv3

#### 6.1. Manual curation

Manual curation of automated predictions was carried out using the Apollo annotation editor [21] plugin for JBrowse genome browser [141]. Among other genomic resources, the Apollo instance for the Diaci v3.0 genome is hosted at citrusgreening.org [1]. All the evidence used for automated annotation was added as tracks on JBrowse/Apollo. Other tracks to assist in accurate evidence-based curation included proteins from related insects mapped with Exonerate [142]. All publicly available RNA-Seq datasets were mapped as quantitative tracks. Manual curation was performed following the workflow previously described in Hosmani *et al.,* [39] and modifications to the workflow, if any, are described in the individual pathway reports. A more detailed version of the annotation workflow is available from Shippy *et al.*, [143].

#### 6.2. Official Gene Set v3 (OGSv3)

The Official gene set version 3 (OGSv3) consists of predicted gene models from the MAKER annotation pipeline and manually curated genes from the Apollo annotation editor. Before merging curated genes to create OGSv3, overlapping genes from the automatic predictions were removed. We also removed 258 predicted genes that were flagged as incorrect by annotators during the manual curation process. These included incorrect predictions, false duplications, and genes not supported by transcriptome evidence.

#### 6.3. Orthology analysis

Orthology analysis of 12 species (Bemisia tabaci, Acyrthosiphon pisum, Rhopalosiphum maidis, Diaphorina citri, Pachypsylla venusta, Drosophila melanogaster, Anopheles gambiae, Homalodisca vitripennis, Gerris buenoi, Cimex lectularius, Halyomorpha halys, Oncopeltus fasciatus) was performed using a total of 240,242 genes. Orthofinder (v2.3.3) [110, 144] was run to get orthogroups using the -S blast parameter as a sequence search program. Orthogroups annotation was performed with Interproscan (v5.32-71.0) [145] to assign GO terms and Pfam domains.

## Declarations

### List of abbreviations

ALT: Alternate
BLAST: Basic Local Alignment Search Tool
*C*Las: “*Candidatus* Liberibacter asiaticus”
DBD: DNA Binding Domains
D. citri: Diaphorina citri, Asian citrus psyllid
FPKM: Fragments per kilobase of exon per million mapped
FTP: File Transfer Protocol
Gb/Mb/Kb/bp: Gigabyte, Megabyte, Kilobyte, base pair
GFF: General Features Format
GO: Gene Ontology
HLB: Huanglongbing or Citrus Greening disease
HQ: High Quality/Low Quality
HSI: Hispanic Serving Institute
OGS: Official Gene Set
ORF: Open Reading Frame
NCBI SRA: National Center for Biotechnological Information Sequence Read Archive
NIFA: National Institute of Food and Agriculture
TF: Transcription Factor
USDA: United States Department of Agriculture
WCGNA: Weighted Gene Co-expression Network Analysis

## Ethics statement

Not applicable.

## Consent for publication

Not applicable.

## Author contributions

SJB, TD, MH, WBH, RGS and LAM conceptualized the project and obtained funding. SJB, TD, SS and LAM contributed to project administration. PSH, MF-G, SS, TDS, MTW, and JB developed methods. PSH, MF-G, and SS curated data and contributed to software development. TDS, SM, MTR, CV, CM, WT, LD, CC, SH, MR, JN, KK, MRJ, DH, SA, YD, TP, SV, RA, SN, JV, MP, AL, STB, NP, YO, BT, RG, AN, DLH, AK, AT, BK, XC, ME, ND, RM, RH, KG, MK, MH, MMS, KMRP, AM, NW, JB, TW, SL, SMi, EM, SK and SS conducted the investigation. TD; SS; TDS; JB, and SJB provided leadership and oversight for the project. TDS, CV, CM, LD, and SM validated manual annotations. MAM, TDS; JB; MTR, and SS created visualizations of the data. PSH; MF-G; TDS; JB; TD; and SS wrote the initial draft. MAM; TDS; WBH, MH and SS reviewed and edited the manuscript. All authors approved the submitted version.

## Acknowledgements

Kascha Bohnenblust Johnson from Kansas State University assisted with the organization of meetings for the community curation effort. Some of the computing for this project was performed on the Beocat Research Cluster at Kansas State University, which is funded in part by NSF grants CNS-1006860, EPS-1006860, EPS-0919443, ACI-1440548, CHE-1726332, and NIH P20GM113109. Maria Gonzalez, Biological Science Technician, USDA, ARS, Ft. Pierce, FL, assisted with psyllid collection, DNA and RNA extractions, and production of high-molecular weight gDNA.

## Disclaimers

Mention of trade names or commercial products herein is solely for the purpose of providing specific information and does not imply recommendation or endorsement, to the exclusion of other similar products or services by the U.S. Department of Agriculture. USDA is an equal opportunity provider and employer.

## Supplementary Information

S0: Github Repo https://github.com/s-hoyt/endosymbiont-analysis Contains scripts, prokka changes logs, unique protein names

S1: Genomes in Custom Database

| **Wolbachia Strains** | **Accession Number** |
| --- | --- |
| Wolbachia endosymbiont of Drosophila melanogaster (a-proteobacteria) | ASM802v1 |
| Wolbachia endosymbiont strain TRS of Brugia malayi (a-proteobacteria) | ASM838v1 |
| Wolbachia endosymbiont of Onchocerca ochengi (a-proteobacteria) | ASM30688v1 |
| Wolbachia endosymbiont of Drosophila simulans wNo (a-proteobacteria) | ASM37658v1 |
| Wolbachia endosymbiont of Cimex lectularius (a-proteobacteria) | ASM82931v1 |
| Wolbachia sp. wRi (a-proteobacteria) | ASM2228v1 |
| Wolbachia endosymbiont of Drosophila simulans wHa (a-proteobacteria) | ASM37660v1 |
| Wolbachia endosymbiont of Onchocerca volvulus str. Cameroon (a-proteobacteria) | W_O_volvulus_Cameroon_v3 |
| Wolbachia endosymbiont of Drosophila simulans wAu (a-proteobacteria) | Wau001 |
| Wolbachia endosymbiont of Diaphorina citri | wACP3 |
| **Carsonella Strains** | **Accession Number** |
| Candidatus Carsonella ruddii PV (g-proteobacteria) | ASM1036v1 |
| Candidatus Carsonella ruddii CE isolate Thao2000 (g-proteobacteria) | ASM28723v1 |
| Candidatus Carsonella ruddii CS isolate Thao2000 (g-proteobacteria) | ASM28725v1 |
| Candidatus Carsonella ruddii HC isolate Thao2000 | ASM28727v1 |
| (g-proteobacteria) |  |
| Candidatus Carsonella ruddii HT isolate Thao2000 (g-proteobacteria) | ASM28729v1 |
| Candidatus Carsonella ruddii PC isolate NHV (g-proteobacteria) | ASM28731v1 |
| Candidatus Carsonella ruddii DC (g-proteobacteria) | ASM44157v1 |
| Candidatus Carsonella ruddii (g-proteobacteria) [YCCR] | ASM127451v1 |
| Candidatus Carsonella ruddii (g-proteobacteria) [BC] | ASM200935v1 |
| **Profftella Strains** | **Accession Number** |
| Candidatus Profftella armatura (b-proteobacteria) | ASM44155v1 |
| Candidatus Profftella armatura (b-proteobacteria) [YCPA] | ASM127878v1 |

S2: Dot Plots and Blast results showing Alignment between our genome assemblies and published genomes

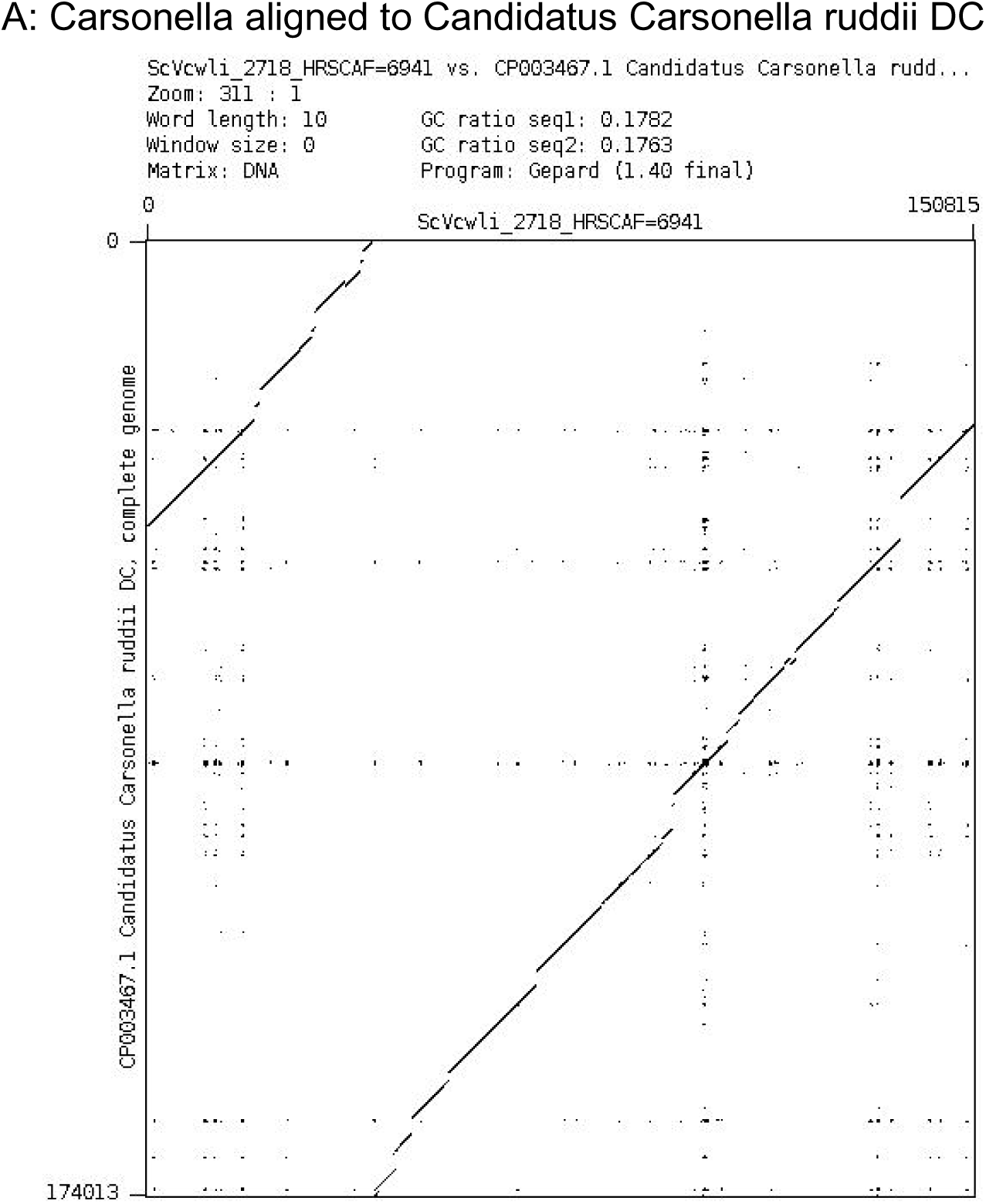
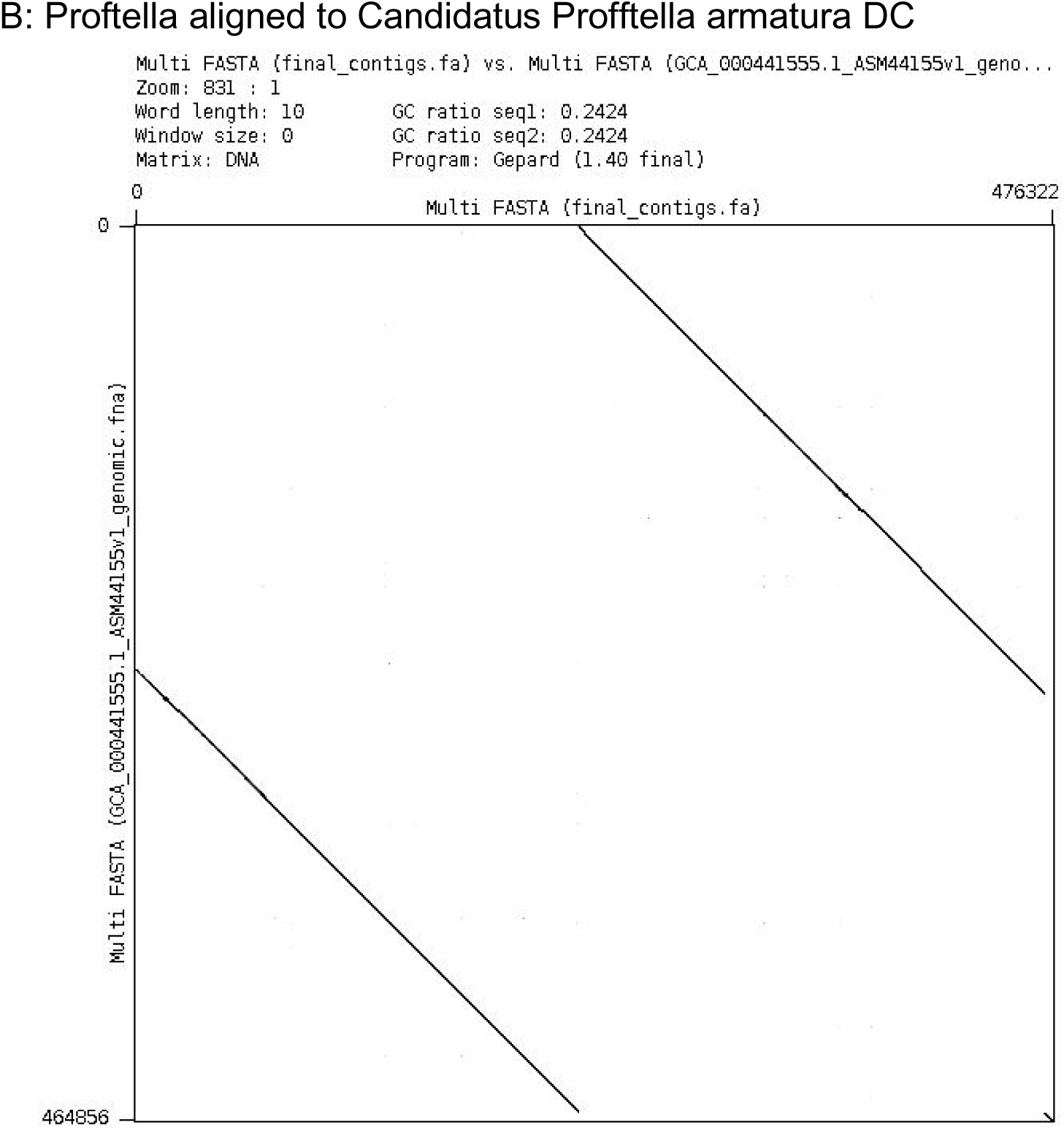
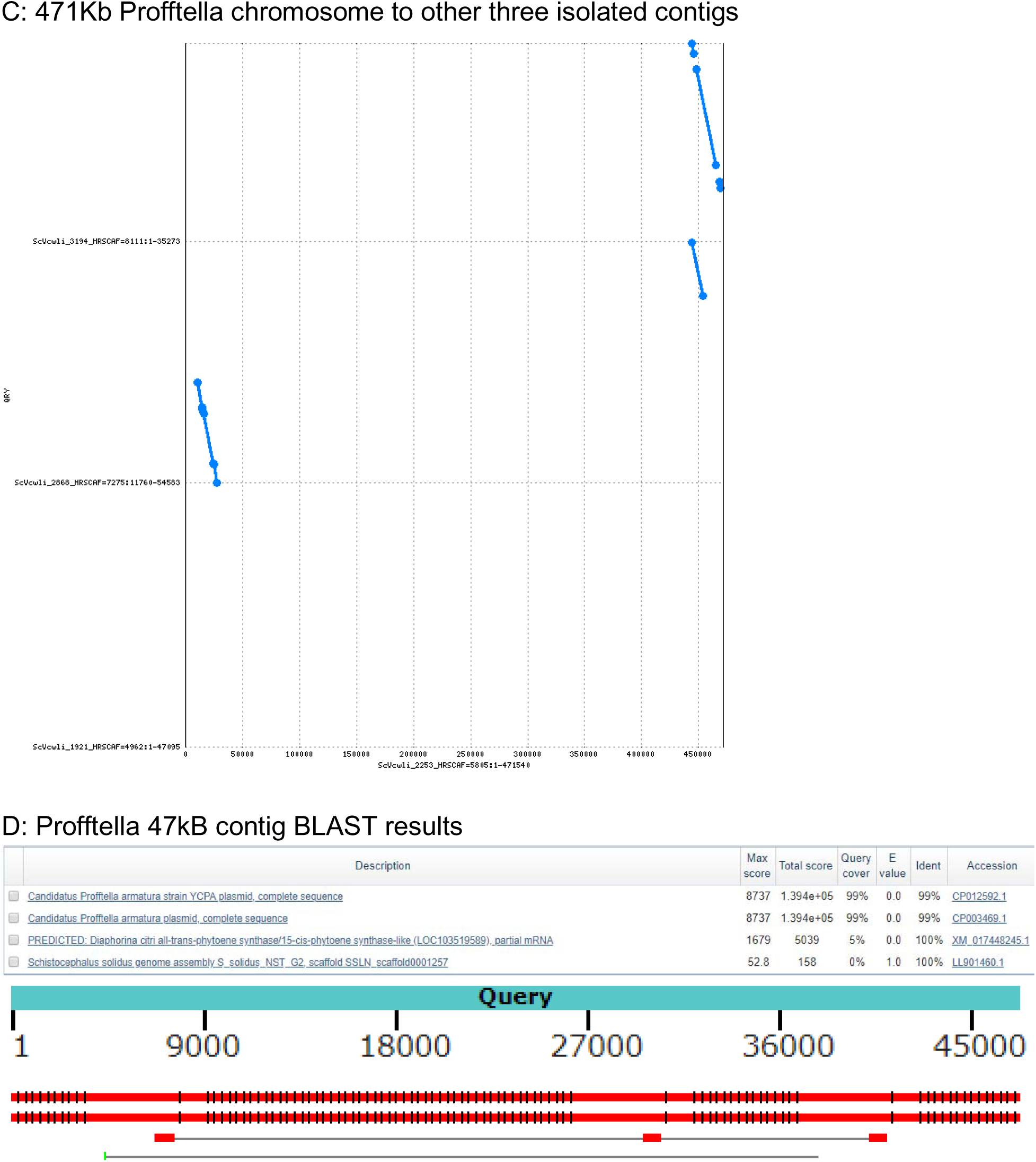
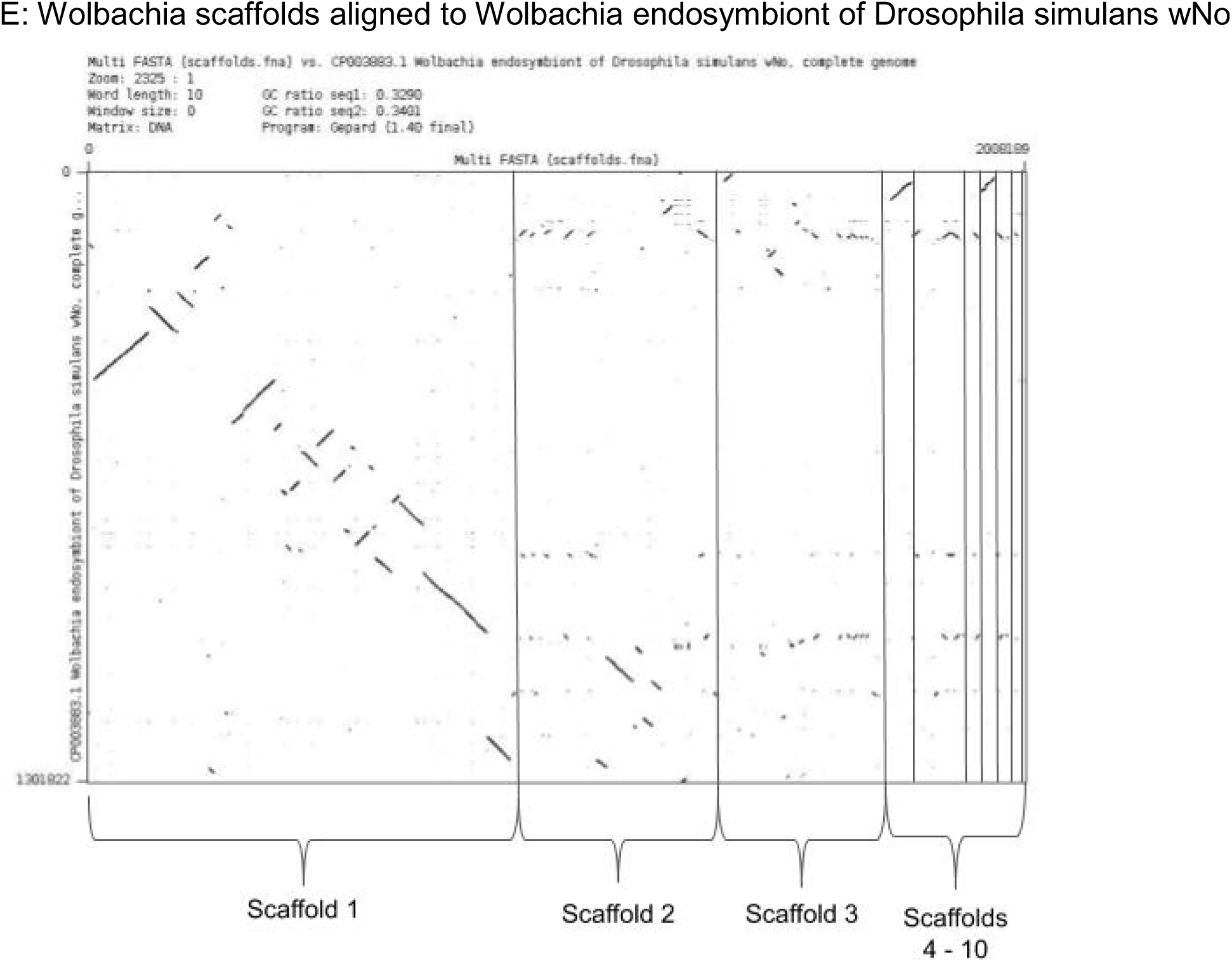

**Supplemental Figure 1:**
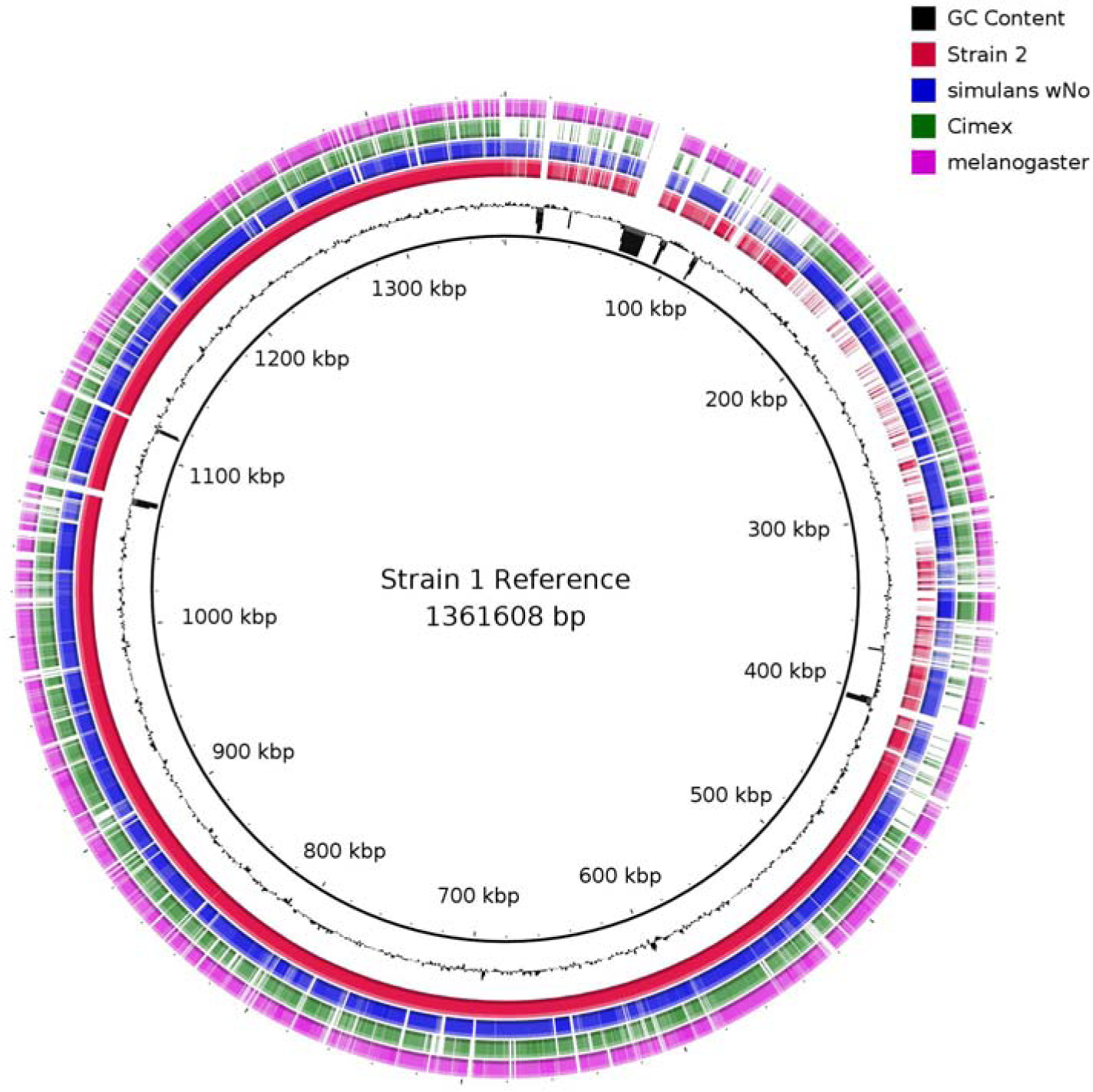
A comparison between the two strains of *Wolbachia* and three published genomes. Regions of dark red indicate where strain 1 and strain 2 share the same sequence; where there is more white indicates gaps where strains 1 and 2 differ. Regions of high GC content indicate regions of the reference genome that do not align to the other genomes at all. melanogaster = *Drosophila melanogaster Wolbachia* symbiont, simulans wNo = *Drosophila simulans Wolbachia* symbiont, cimex = *Wolbachia* symbiont of *Cimex lectularis*. Accession numbers and references for all *Wolbachia*, *Carsonella* and *Profftella* genomes used for comparison to *D. citri* endosymbionts are shown in the table.

| Wolbachia Strains | Accession Number | Reference |
| --- | --- | --- |
| Wolbachia endosymbiont of *Drosophila melanogaster* (α-proteobacteria) | ASM802v1 | (Wu et al. 2004) |
| Wolbachia endosymbiont strain TRS of *Brugia malayi* (a-proteobacteria) | ASM838v1 | (Foster et al. 2005) |
| Wolbachia endosymbiont of *Onchocerca ochengi* (a-proteobacteria) | ASM30688v1 | (Darby et al. 2012) |
| Wolbachia endosymbiont of *Drosophila simulans* wNo (a-proteobacteria) | ASM37658v1 | (Ellegaard et al. 2013) |
| Wolbachia endosymbiont of *Cimex lectularius* (a-proteobacteria) | ASM82931v1 | (Nikoh et al. 2014) |
| Wolbachia sp. wRi (a-proteobacteria) | ASM2228v1 | (Klasson et al. 2009) |
| Wolbachia endosymbiont of *Drosophila simulans* wHa (a-proteobacteria) | ASM37660v1 | (Ellegaard et al. 2013) |
| Wolbachia endosymbiont of *Onchocerca volvulus* str. Cameroon (a-proteobacteria) | W_O_volvulus_Cameroon_v3 | No publication listed* |
| Wolbachia endosymbiont of *Drosophila simulans* wAu (a-proteobacteria) | Wau001 | (Sutton et al. 2014)** |
| Wolbachia endosymbiont of *Diaphorina citri* | wACP3 | (Saha et al. 2012)^ |
| **Carsonella Strains** | **Accession Number** |  |
| Candidatus *Carsonella ruddii* PV (g-proteobacteria) | ASM1036v1 | (Nakabachi et al. 2006) |
| Candidatus *Carsonella ruddii* CE isolate Thao2000 (g-proteobacteria) | ASM28723v1 | (Sloan and Moran 2012) |
| Candidatus *Carsonella ruddii* CS isolate Thao2000 (g-proteobacteria) | ASM28725v1 |  |
| Candidatus *Carsonella ruddii* HC isolate Thao2000 (g-proteobacteria) | ASM28727v1 |  |
| Candidatus *Carsonella ruddii* HT isolate Thao2000 (g-proteobacteria) | ASM28729v1 |  |
| Candidatus Carsonella ruddii PC isolate NHV (g-proteobacteria) | ASM28731v1 |  |
| Candidatus Carsonella ruddii DC (g-proteobacteria) | ASM44157v1 | (Nakabachi et al. 2013) |
| Candidatus Carsonella ruddii (g-proteobacteria) [YCCR] | ASM127451v1 | (Wu et al. 2015) |
| Candidatus Carsonella ruddii (g-proteobacteria) [BC] | ASM200935v1 | No publication listed^^ |
| **Profftella Strains** | **Accession Number** |  |
| Candidatus Profftella armatura (b-proteobacteria) | ASM44155v1 | (Nakabachi et al. 2013) |
| Candidatus Profftella armatura (b-proteobacteria) [YCPA] | ASM127878v1 | (Wu et al. 2015) |

\*No publication found, but can find the data using this assembly ID and BioProject PRJEB4840 or GenBank GCA\_000530755.1
\*\*Can find the data using this assembly ID and BioProject PRJEB6321 or GenBank GCA\_000953315.1
^Can find the data using this assembly ID. Could use BioProject PRJNA29451 or GenBank GCA\_000331595.1
^^No publication or accession found

**Supplemental Table 1:**
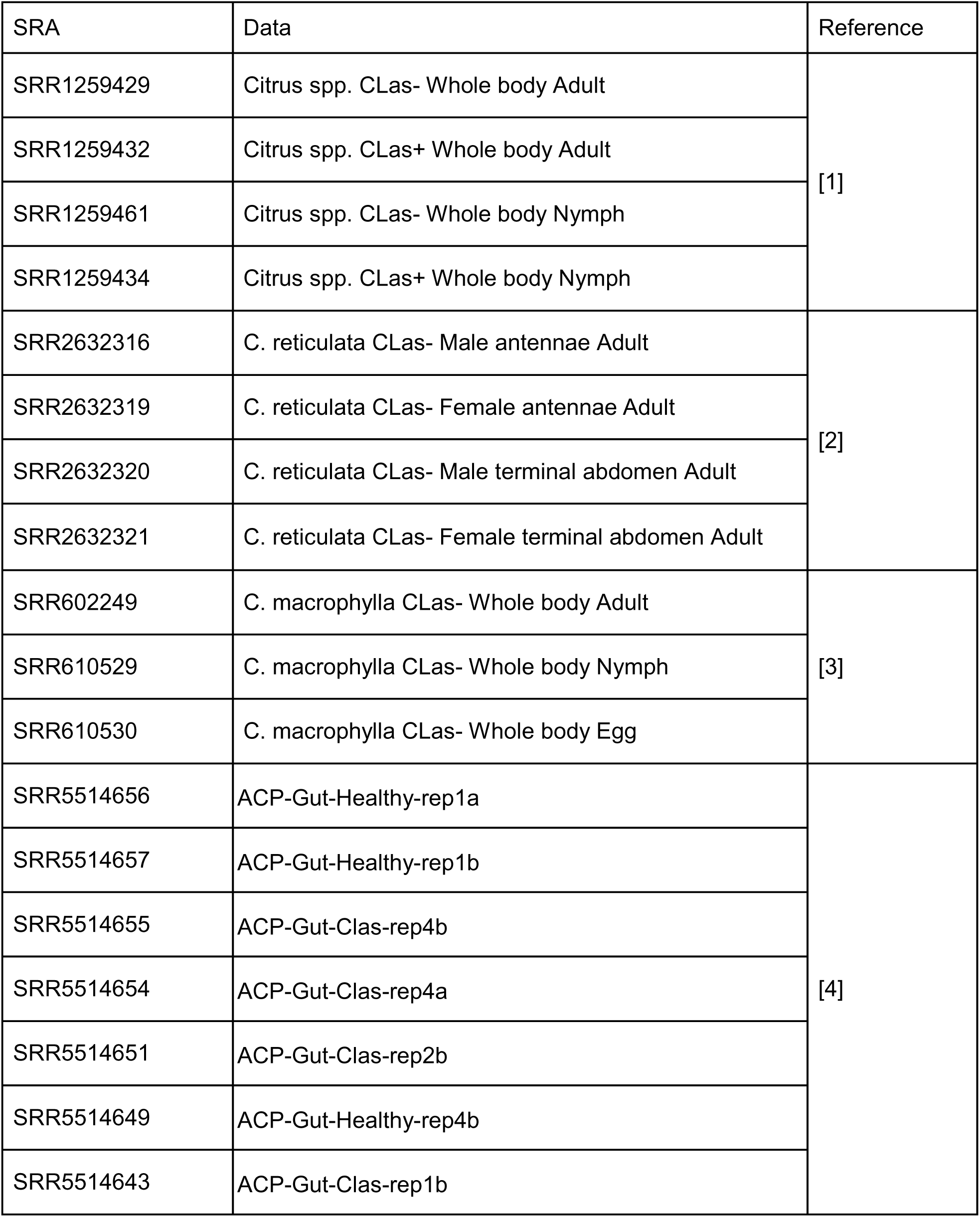
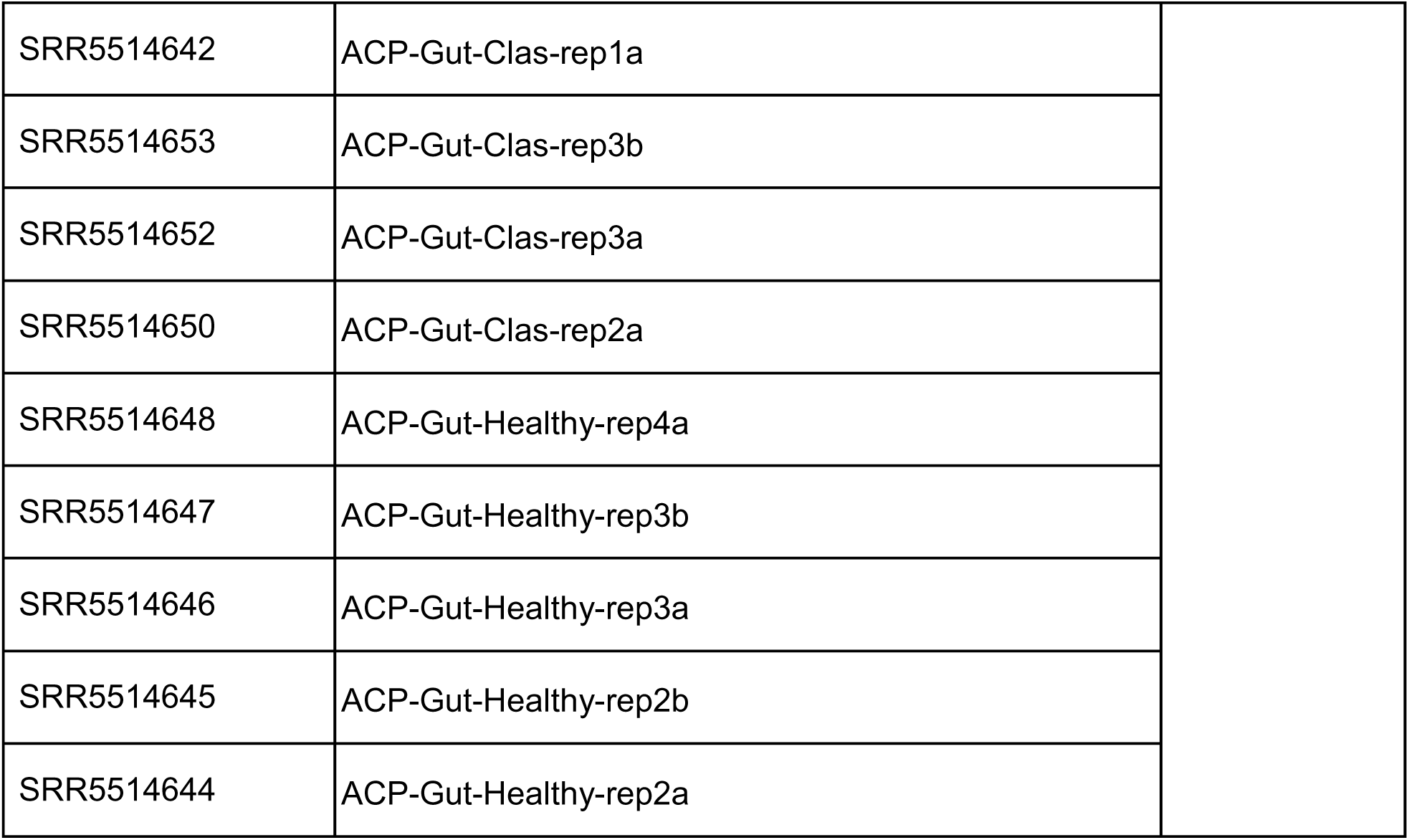
RNA-seq data used for genome annotation and de novo transcriptome assembly.

---

**Supplemental Table 2:** BUSCO statistics. Percentages of the Arthropoda (arthropoda_odb10) and Hemiptera (hemiptera_odb10) BUSCO gene set that are complete, duplicated, fragmented or missing in Florida *Diaphorina. citri* genome versions, official gene sets and transcriptomes.

|  | BUSCO Arthropoda 1013 Gene Set (%) |  |  |  |
| --- | --- | --- | --- | --- |
|  | Complete | Duplicated | Fragmented | Missing |
| **Diaci v3.0** | 93.5 | 26.4 | 3.3 | 3.2 |
| **Diaci v2.0** | 92.1 | 36.8 | 4.6 | 3.3 |
| **Diaci v1.1** | 82.6 | 4.6 | 11.4 | 6.0 |
| ***de novo* transcriptome** | 88 | 30.8 | 3.5 | 8.5 |

|  | BUSCO Hemiptera 2510 Gene Set (%) |  |  |  |
| --- | --- | --- | --- | --- |
|  | Complete | Duplicated | Fragmented | Missing |
| **Diaci v3.0** | 93.8 | 33.6 | 3.3 | 2.9 |
| **Diaci v2.0** | 93.8 | 33.6 | 3.3 | 2.9 |
| **Diaci v1.1** | 88.9 | 4.7 | 7.0 | 4.1 |
| ***de novo* transcriptome** | 94.2 | 24 | 2.6 | 3.2 |

